# P-body sequestration of clock transcripts delays repressor synthesis to set circadian period in *Drosophila*

**DOI:** 10.64898/2026.09.28.754909

**Authors:** Yangbo Xiao, Ye Yuan, Shrabastee Chakraborty, David Brooks, Swathi Yadlapalli

**Affiliations:** Department of Cell and Developmental Biology, University of Michigan, Ann Arbor, MI; Michigan Neuroscience Institute, University of Michigan, Ann Arbor, MI

## Abstract

Negative-feedback oscillators require a delay between the accumulation of a repressor’s mRNA and the action of its protein. In the circadian clock, this delay has been attributed largely to post-translational control of PERIOD (PER) stability and nuclear entry. The RNA-binding proteins shown to regulate *per* translation, ATAXIN-2 and its partners, promote it, leaving open whether any step holds clock transcripts back before they are translated. Here, using time-resolved miniTurbo proximity labeling of endogenous PER across four phases of the circadian cycle in *Drosophila* clock neurons, we define a 252-protein PER proximitome that partitions into a nuclear arm and a cytoplasmic RNA-metabolism arm. A behavioral RNAi screen identified two P-body components, the DEAD-box helicase Me31B (DDX6) and the 5′–3′ exonuclease Pacman (Pcm; XRN1), as strong regulators of circadian rhythms. Using single-molecule RNA-FISH, proximity RNA editing and ribosome profiling, we show that as *per* and *tim* transcripts accumulate, they localize to Me31B-labeled P-bodies and are poorly translated, most prominently at ZT12. Me31B knockdown disrupts P-bodies and releases *per* mRNA from them, causing PER to accumulate earlier and to ∼2-fold higher levels, whereas Me31B overexpression delays PER accumulation and lengthens the free-running period by ∼2 h. Knockdown of Pcm, in contrast, impairs clearance of *per* mRNA, sustaining PER and TIM accumulation, prolonging the repression phase and abolishing cycling of ∼89% of rhythmic transcripts. Together, these findings identify P-body sequestration as a repressive step that delays repressor synthesis, and Pcm-dependent decay as required to end repression on time. Given the deep conservation of DDX6 and XRN1, RNP compartments may provide a conserved means of generating delay in circadian and other negative-feedback circuits.

## Introduction

Animals coordinate behavior, physiology, and metabolism with the 24-hour day through cell-autonomous circadian clocks. In *Drosophila* and mammals, these clocks are based on transcription–translation feedback loops (TTFLs), in which transcriptional activators (CLK/CYC in flies; CLOCK/BMAL1 in mammals) drive cyclic expression of repressor genes (*per*/*tim* in flies; *Per*/*Cry* in mammals) that feed back to terminate their own transcription^1, 2^. To prevent rapid dampening, sustained ∼24-hour oscillations require a multi-hour interval between repressor mRNA accumulation and protein accumulation^3–6^. In *Drosophila*, *per* and *tim* mRNAs peak in the early night^7^, whereas PER and TIM proteins peak several hours later^8, 9^. While nuclear PER regulation has been mapped at high resolution^10–13^, the cytoplasmic machinery that shapes this interval remains poorly characterized.

Known cytoplasmic clock regulators act primarily post-translationally or by altering transcript abundance. Kinases such as DOUBLETIME and CASEIN KINASE 2 phosphorylate PER to gate its nuclear entry^14, 15^, priming it for SLIMB-mediated degradation^16–18^. At the mRNA level, factors like ATX2 promote translational activation of *per*^19–21^, while microRNAs and the POP2 deadenylase tune transcript stability^22–25^. Although the helicase Me31B/DDX6 was previously implicated in circadian behavior, it was reported to function independently of PER translation^26^. Consequently, whether clock transcripts are held in a translationally silent state to delay repressor synthesis, and what machinery would execute such a step, remains unknown.

To identify cytoplasmic regulators of PER without a candidate bias, we set out to map the neighborhood of PER itself. This has been difficult because core clock proteins form dynamic, membrane-less condensates^9, 27^, and standard biochemical extraction can dissolve phase-separated structures and fails to retain short-lived, low-affinity interactions^28, 29^. Proximity-dependent biotinylation circumvents these limitations: the engineered ligase miniTurbo covalently labels spatial neighbors within ∼10 nm of a target in intact, living cells, capturing both stable and transient associations within defined temporal windows^30, 31^. Here, we apply miniTurbo proximity labeling of endogenous PER across the clock neurons of the adult *Drosophila* brain at six-hour resolution to map the temporal PER proximitome. The dataset partitions into a nuclear arm enriched for transcriptional regulators and a cytoplasmic arm enriched for mRNA-binding and mRNA-decay factors. In a behavioral RNAi screen of PER-proximal candidates, the two strongest hits converged on a single compartment: the DEAD-box helicase Me31B (DDX6 ortholog) and the 5′–3′ exonuclease Pacman (Pcm; XRN1 ortholog), both canonical components of cytoplasmic processing bodies (P-bodies)^32, 33^.

Notably, P-bodies are the compartment where translational repression and transcript decay converge. They serve as reversible storage depots for translationally silenced but intact transcripts, which can exit and return to polysomes^34–37^, and they also concentrate the core decay machinery^33, 38^; whether a given transcript is stored or degraded remains an open question in the field^32^. Me31B is a well-established repressor of stored mRNAs, silencing transcripts in both the germline^39, 40^ and neurons^41^. Its mammalian ortholog, DDX6, similarly represses translation through interactions with the CCR4–NOT complex and the eIF4E-binding protein 4E-T^42–44^. Pcm degrades decapped transcripts in the 5′-to-3′ direction^45, 46^. Whether P-bodies sequester clock transcripts, and whether Pcm clears them, have not been tested.

Using single-molecule RNA-FISH, proximity RNA editing, ribosome profiling, and bidirectional genetic perturbation, we find that *per* and *tim* transcripts associate with Me31B-labeled P-bodies as they accumulate, most prominently at ZT12, when they are poorly translated. Me31B knockdown disrupts P-bodies, releases *per* mRNA, and causes PER to accumulate earlier and to ∼2-fold higher levels, whereas overexpression of Me31B delays PER accumulation and lengthens the free-running period by ∼2 h. Knockdown of Pcm leaves *per* mRNA elevated at its trough, sustains nuclear PER and TIM, prolongs the repression phase, and abolishes cycling of ∼89% of rhythmic transcripts. These results identify P-body sequestration as a step that holds clock transcripts in a translationally silent state to delay repressor synthesis, and Pcm-dependent decay as required to end repression on time.

## Results

### Time-resolved proximity labeling of endogenous PER defines a 252-protein proximitome in the adult brain

To map PER’s protein neighborhood across the circadian cycle *in vivo*, we generated a knock-in allele fusing miniTurbo (mTb) to the C-terminus of endogenous PER (*per-V5-mTb*; **Figure 1A**). As a matched negative control, we used a *per-mNG* knock-in carrying mNeonGreen at the identical genomic position^9^, which accounts for endogenously biotinylated proteins and non-specific matrix binders. We next tested whether the tagged allele is functional. In 12:12 light:dark conditions, *per-V5-mTb* flies were behaviorally indistinguishable from controls, and in constant darkness they free-ran with a period of ∼24 h comparable to wild-type controls assayed in parallel (**Figure 1B**). The allele also restored rhythmicity to *per ¹* null mutants, with *per-V5-mTb*/*per ¹* females free-running at ∼25 h; this modest period lengthening is likely attributable to a gene dosage effect from a single functional copy of *per* (**Figure 1C**). Finally, anti-V5 immunoblots on head extracts collected every 4 h showed that PER-V5-mTb abundance oscillated across the 24-h cycle, peaking near ZT20 and reaching a trough near ZT8 (**Ext. Data Figure 1A**), matching the reported profile of endogenous PER. Together, these results indicate that PER-V5-mTb is functional and retains normal temporal regulation.

**Fig. 1.**
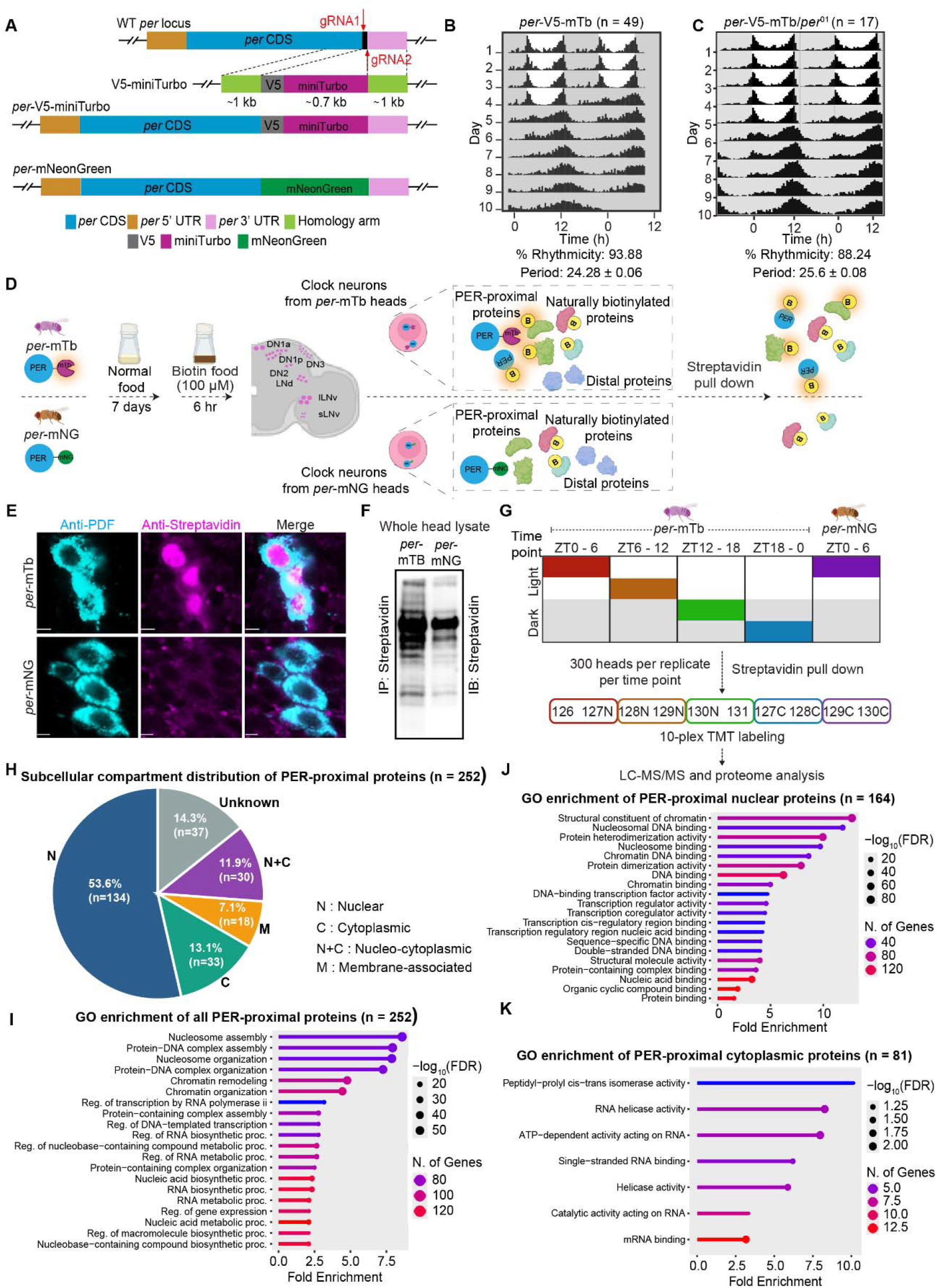
A time-resolved proximity proteome of PER. **(A)** CRISPR/Cas9 strategy for generating *per*-V5-miniTurbo flies, with V5 and miniTurbo fused in frame to the C-terminus of endogenous PER (top), and the *per*-mNeonGreen knock-in used as the no-ligase control (bottom). Guide RNA positions and 1 kb homology arms are indicated. **(B, C)** Double-plotted population actograms of *per*-V5-miniTurbo (n = 49) (B) and *per*-V5-miniTurbo/*per ¹* (n = 17) (C) flies entrained to 12:12 LD (ZT0, lights on; ZT12, lights off) and released into constant darkness. Percent rhythmicity and free-running period (mean ± SEM) are shown below. **(D)** Labeling strategy. Flies were fed 100 µM biotin for a 6-h window, during which miniTurbo biotinylates proteins near PER in clock neurons; biotinylated proteins were captured from head lysates on streptavidin beads. **(E)** Whole-mount brains stained for PDF (cyan) and biotinylated proteins (streptavidin, magenta), showing large ventral lateral neurons. Scale bars, 5 µm. **(F)** Streptavidin blot of head lysates following streptavidin pull-down. **(G)** TMT 10-plex design. *per*-V5-miniTurbo samples were labeled during one of four 6-h windows (ZT0–6, ZT6–12, ZT12–18, ZT18–24) and *per*-mNeonGreen controls at ZT0–6; ∼300 heads per replicate, two biological replicates per condition. TMT channel assignments are shown. **(H)** Subcellular compartment distribution of the 252 PER-proximal proteins (log FC ≥ 0.5 and FDR-adjusted p < 0.05 in at least one matched window). **(I–K)** GO enrichment of all (n = 252) (I), nuclear (n = 164) (J) and cytoplasmic (n = 81) (K) PER-proximal proteins. Dot size, −log (FDR); color, number of genes.mTb, miniTurbo; mNG, mNeonGreen; IP, immunoprecipitation; IB, immunoblot; PDF, pigment-dispersing factor; ZT, Zeitgeber time; FC, fold change; FDR, false discovery rate.

For proximity profiling, 7-day-old *per-V5-mTb* or *per-mNG* adults entrained to 12:12 light:dark cycles were transferred to food supplemented with 100 µM biotin for 6-h labeling windows timed to one of four Zeitgeber phases (ZT0–6, 6–12, 12–18, 18–24). Following labeling, fly heads were collected, lysed, and biotinylated proteins were captured on streptavidin beads (**Figure 1D**). Immunofluorescence staining of whole-mount brains with anti-streptavidin and anti-PDF revealed robust, ligase-dependent labeling in PDF-expressing clock neurons in *per-V5-mTb* flies, whereas *per-mNG* controls showed only background signal (**Figure 1E**). Blots probed with streptavidin-HRP showed extensive biotinylation in *per-V5-mTb* extracts and negligible signal in *per-mNG* controls (**Figure 1F, Ext. Data Figure 1B**), indicating that capture reflects ligase activity rather than PER abundance. Across all four labeling windows, streptavidin eluates from *per-V5-mTb* heads yielded numerous biotinylated species that was reduced to a sparse background of endogenously biotinylated proteins in *per-mNG* controls (**Ext. Data Figure 1B**). Because these endogenously biotinylated background proteins (predominantly mitochondrial carboxylases) are recovered in both genotypes and are therefore subtracted in our ratiometric (*per-V5-mTb* / *per-mNG*) quantitative analysis (see Methods).

We tiled four non-overlapping 6-h windows in *per-mTb* flies (ZT0-6, 6-12, 12-18, and 18-0), chosen to span the phases of nuclear PER degradation, the cytoplasmic trough, cytoplasmic PER accumulation prior to nuclear translocation, and peak nuclear PER (**Figure 1G**). This design tracks PER’s subcellular distribution across the day, established in fixed tissue and directly visualized in live clock neurons using fluorescent knock-in alleles^9^. A *per-mNG* sample collected at ZT0–6 served as the shared denominator for all *per-mTb* windows; endogenously biotinylated proteins were recovered at comparable intensities across the four *per-V5-mTb* windows (**Ext. Table 1**), indicating that overall labeling efficiency did not vary systematically with time of day and supporting the use of a single reference sample. All ten samples (two biological replicates, five conditions, ∼300 fly heads each, ∼3,000 heads in total) were on-bead trypsinized, TMT 10-plex labeled, pooled, and analyzed in a single LC-MS/MS run, avoiding between-run variation across the time course. Replicate-pair correlations were high (*R*^2^ = 0.78 - 0.97; **Ext. Data Figures 1C-1G**). After primary MS-quality filtering (≥ 2 unique peptides, valid quantification across all four windows, and gene-level isoform collapse), we removed proteins belonging to categories that are abundant carry-over contaminants in fly head preparations (yolk/fat-body and extracellular matrix proteins) and are recovered equivalently in both genotypes; none of the high-confidence hits reported below came from these categories. The remaining 2,969 proteins formed the background proteome.

For each protein, we computed log_2_ (*per*-mTb/ *per*-mNG) at each window, together with the corresponding Benjamini-Hochberg-adjusted *p*-value, which estimates variance across proteins and is appropriate for two biological replicates. Proteins were classified as PER-proximal at log_2_ FC ≥ 0.5 (≥1.41-fold enrichment over *per-mNG*) and adj *p* < 0.05 in at least one window, yielding 252 high-confidence hits. The compartment- and pathway-level enrichments described below remained stable across more stringent thresholds (log_2_ FC ≥ 0.7 and ≥ 1.0), indicating that they are not sensitive to the specific cutoff.

The 252-protein set was strongly nuclear-biased. Using GO cellular component / UniProt annotations, we assigned hits to nuclear (n = 164) and cytoplasmic (n = 81) subsets, with nucleo-cytoplasmic shuttling proteins assigned to both (**Figure 1H**). This distribution is consistent with PER’s established subcellular localization, supporting the spatial specificity of in vivo proximity labeling. GO Biological Process enrichment of the full dataset centered on chromatin organization and regulation of gene expression (**Figure 1I**). Compartment-stratified GO analysis recovered two complementary signatures: nuclear hits were dominated by chromatin and nucleosomal DNA binding, transcription regulator activity, and chromatin remodeling (**Figure 1J**), while cytoplasmic hits were enriched for peptidyl-prolyl cis–trans isomerase activity, RNA helicase activity, and mRNA binding (**Figure 1K**). The cytoplasmic signature was less significant than its nuclear counterpart, consistent with the smaller number of cytoplasmic hits and with PER’s nuclear-biased residence time. Nonetheless, two of the recovered terms, RNA helicase activity and mRNA binding, pointed toward post-transcriptional regulation; the peptidyl-prolyl isomerase signature is a separate lead that we did not pursue here.

KEGG and Reactome analysis identified enrichment for basal transcription factors (BH-adjusted p < 10), histone modification and chromatin remodeling, spliceosome, and RNA degradation (**Ext. Data Figure 1H**). The circadian rhythm pathway was also recovered, an internal validation that the proximitome captures the biology of the bait. By contrast, generic signaling pathways such as Wnt and Notch signaling, as well as endocytosis were not enriched, indicating that the 252-protein set reflects a focused PER-proximal network rather than abundant cellular machinery. Together, these data establish a time-resolved proximity-labeling system for endogenous PER in the adult brain and define a 252-protein proximitome that partitions into a nuclear chromatin/transcription arm and a cytoplasmic RNA-metabolism arm.

### The PER proximitome resolves into phase-specific nuclear and cytoplasmic modules

A four-way comparison of PER-proximal proteins across windows showed that 134 of 252 (53.2%) were significant in a single window, 99 (39.3%) in two, typically adjacent, windows, 8 (3.2%) in three, and 11 (4.4%) in all four (**Figures 2A** and **2B**). Volcano plots revealed phase-specific hit sets (**Figures 2C–2F**). At ZT0–6, the enriched set included core clock components (PER itself, as expected for the bait, along with CYC and CLK), the H3K9 demethylase KDM4B, the Polycomb-associated factors JARID2 and E(PC), the transcriptional corepressor GUG, and the cyclophilin MOCA-CYP (**Figure 2C**). At ZT6–12, hits included TIM and PER, the cytoplasmic regulator LIG, the phosphatase PTPA, the kinase regulator CKIIβ, the chaperone DNAJ-H, and the mRNA-decay factors PCM and the decapping activator EDC3, which we return to below (**Figure 2D**). At ZT12–18, additional translation and RNA-binding proteins appeared (TYF, AGO2, CHMP1, DDX1) (**Figure 2E**). At ZT18–24, the enriched set reorganized markedly to include nuclear pore components (NUP50, NUP214), the Mediator subunit MED8, the splicing factor SF3B3, the hnRNP HRB87F, multiple silencers (SIRT1, SIN3A, E(PC), KDM4B), SWI/SNF subunits (BAP111, MOR), the kinase DCO (DOUBLETIME), the corepressor BON, and the clock effector PDP1 (**Figure 2F**).

**Fig. 2.**
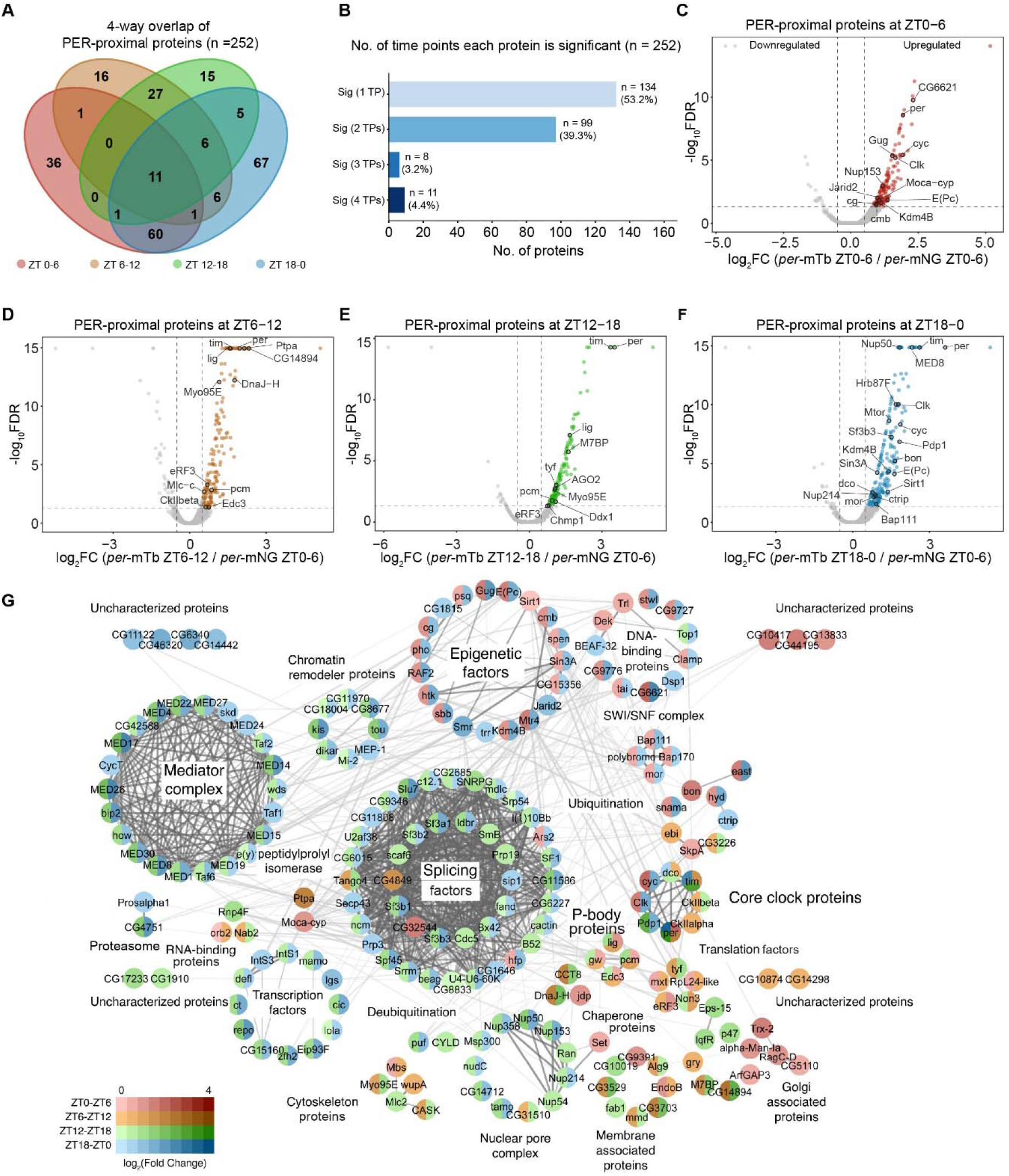
The PER proximitome organizes into distinct nuclear and cytoplasmic modules across circadian phases. **(A)** Four-way Venn diagram of PER-proximal proteins (n = 252), showing proteins significant at a single window or shared across windows. **(B)** Number of PER-proximal proteins significant at one, two, three or all four windows. **(C–F)** Volcano plots of PER-proximal proteins at ZT0–6 (C), ZT6–12 (D), ZT12–18 (E) and ZT18–24 (F). Each window is plotted as log FC of *per*-V5-mTb at that window over the *per*-mNeonGreen control at ZT0–6. Colored points, proteins passing log FC ≥ 0.5 and FDR-adjusted p < 0.05; selected hits are labeled. Dashed lines mark the fold-change and significance thresholds. **(G)** STRING network of all 252 PER-proximal proteins. Nodes are colored by log FC at each of the four windows (color key, lower left); edges represent STRING interaction confidence. Clusters are annotated by functional category. ZT, Zeitgeber time; mTb, miniTurbo; mNG, mNeonGreen; FC, fold change; FDR, false discovery rate.

A STRING-based network of all 252 proteins, with each node colored by log_2_ FC across the four windows, resolved into discrete clusters whose nodes were almost entirely populated by hits from one or two adjacent windows (**Figure 2G**). Major clusters included Mediator, the spliceosome, peptidyl-prolyl isomerases, SWI/SNF and chromatin remodelers, epigenetic factors, the core clock, the nuclear pore, P-body and mRNA-decay factors, and translation factors. Nuclear and cytoplasmic PER are proximal to largely non-overlapping multiprotein neighborhoods.

The nuclear module included components of the nuclear pore complex at ZT18–24, specifically nucleoporins of the nuclear basket and cytoplasmic filaments (NUP50, NUP358, NUP214) (**Ext. Data Figure 2A**). PER’s nuclear neighborhood also shifted with phase.

Components of the general transcription machinery, including Mediator, TFIID and SWI/SNF, were enriched at ZT18–24 (**Ext. Data Figure 2B**), consistent with PER residing near promoter-bound transcriptional machinery during the repression phase. By contrast, a core set of chromatin silencers (CTBP, SIN3A, E(PC), JARID2) was enriched in both nuclear windows (**Ext. Data Figure 2C**). The nuclear proximitome therefore distinguishes phase-restricted association with promoter-bound machinery from sustained association with chromatin silencers across the repression phase. The cytoplasmic module is dominated by mRNA-binding and mRNA-decay factors and includes six canonical P-body components (PCM, ME31B, EDC3, PATR-1, LSM3, GE-1) (**Figures 1K, 2D,** and **2G**), a class not previously linked to core clock regulation.

### P-body components are required for circadian rhythmicity and lie adjacent to PER cytoplasmic foci

To ask whether these proximitome hits reflect functional dependencies of the clock, we performed a behavioral RNAi screen. Using the pan-clock-neuron driver *tim*-UAS-GAL4 (TUG), we knocked down 61 PER-proximal candidates and scored locomotor rhythmicity and free-running period in constant darkness (**Figures 3A, 3B,** and **Ext. Data Figure 3, Table S2**). The screen identified rhythm-disrupting hits across several categories, including ubiquitin-proteasome components (HYD, PROSβ5, USP5) and transcriptional regulators (SMR, LOLA), as well as period-lengthening knockdowns (SIRT1, PUF). The two most severely arrhythmic lines, however, were both P-body components: TUG>*me31B*-RNAi reduced the fraction of rhythmic flies to 8.5% and TUG>*pcm*-RNAi to 10.6%, compared with 86.4% and 94.7% in the corresponding heterozygous RNAi controls and 100% in TUG>+ alone (**Figures 3C–3F** and **Ext. Data Figure 3A, Table S2**). ME31B and PCM act at opposite ends of the mRNA life cycle: the DEAD-box helicase ME31B represses translation of P-body-resident transcripts, whereas the 5′–3′ exonuclease Pacman degrades decapped mRNAs. That an unbiased screen converged on two proteins from the same compartment, acting at these distinct steps, implicates P-body-associated RNA regulation in clock function.

**Fig. 3.**
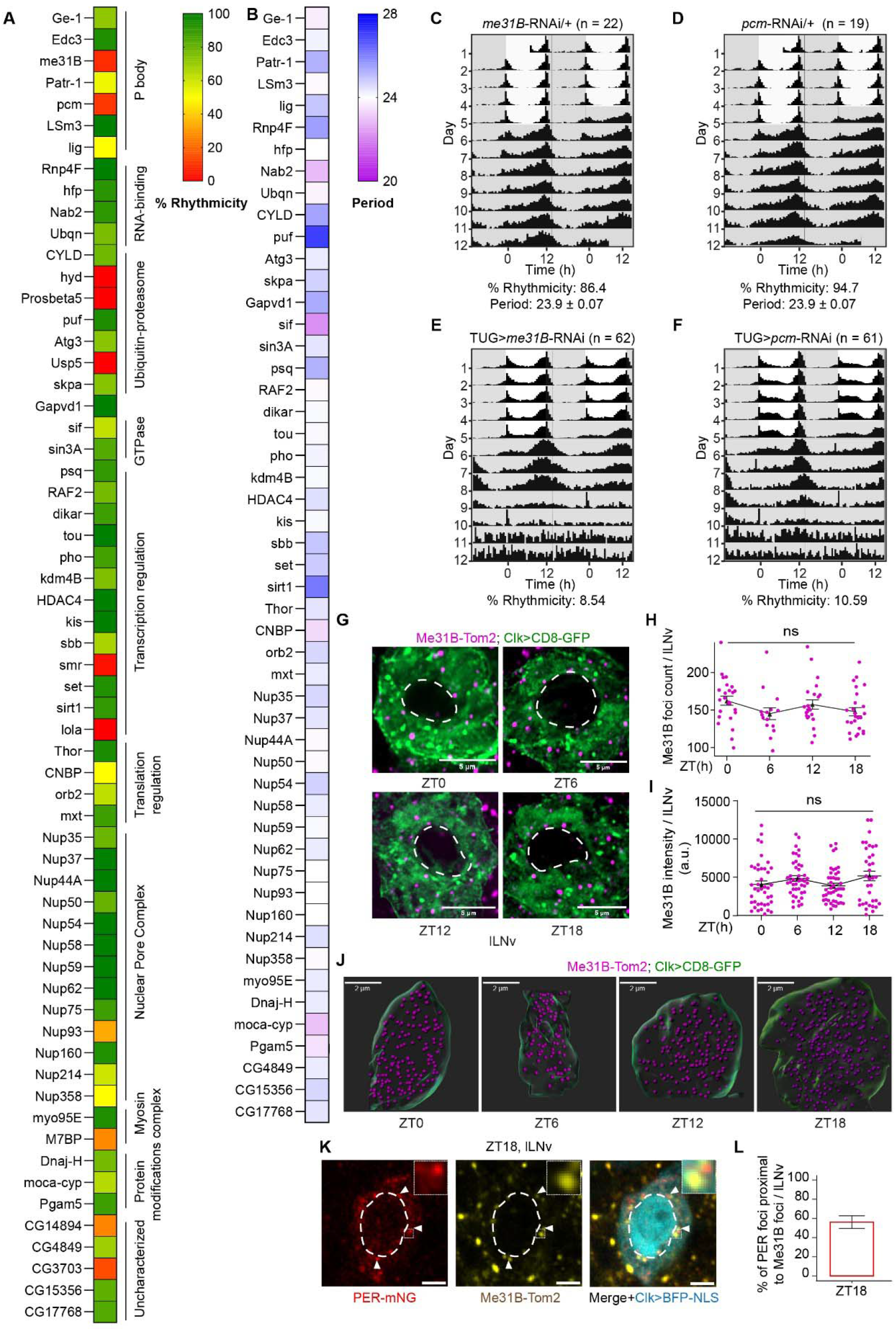
P-body components are required for circadian rhythmicity and lie adjacent to PER cytoplasmic foci. **(A)** Heat map of percent rhythmicity from the RNAi screen of PER-proximal candidates driven by *tim*-UAS-GAL4 (TUG). Candidates are grouped by functional category. **(B)** Free-running period for the same screen; only candidates with rhythmicity > 30% were included. **(C–F)** Double-plotted population actograms of *me31B*-RNAi/+ (n = 22) (C), *pcm*-RNAi/+ (n = 19) (D), TUG > *me31B*-RNAi (n = 62) (E) and TUG > *pcm*-RNAi (n = 61) (F) flies entrained to 12:12 LD (ZT0, lights on; ZT12, lights off) and released into constant darkness. Percent rhythmicity and, where flies were rhythmic, free-running period (mean ± SEM) are shown below. **(G)** Representative images of large ventral lateral neurons (lLNv) in whole-mount brains of *Clk*>CD8-GFP; Me31B-Tom2 flies at ZT0, ZT6, ZT12 and ZT18, showing Me31B-Tom2 foci (magenta) and the CD8-GFP membrane marker (green). Dashed lines demarcate nuclei. Scale bars, 5 µm. **(H, I)** Me31B foci count (H) and integrated intensity (I) per lLNv across the four time points. **(J)** Representative 3D renderings of lLNv at each time point, with segmented Me31B puncta shown as magenta spheres. Scale bars, 2 µm. **(K)** Representative images of lLNv in whole-mount brains of *Clk*>BFP-NLS; *per*-mNeonGreen; Me31B-Tom2 flies at ZT18, showing PER-mNeonGreen (red), Me31B-Tom2 (yellow) and BFP-NLS (cyan). Arrowheads mark PER foci adjacent to Me31B foci; insets show magnified examples. Bright cyan marks the nucleus; faint cytoplasmic cyan reflects the site of BFP-NLS translation. Dashed lines demarcate nuclei. Scale bars, 5 µm. **(L)** Percentage of PER-mNeonGreen foci associated with Me31B foci at ZT18, quantified from (K). Each data point represents a single neuron. TUG, *tim*-UAS-GAL4; lLNv, large ventral lateral neuron; ZT, Zeitgeber time; ns, not significant; a.u., arbitrary units. Statistical tests and exact p-values are given in Table S3.

To test whether the proximity-labeling signal reflects a spatial relationship between PER and P-bodies in clock-neuron cytoplasm, we first visualized endogenous P-bodies using a Me31B-Tom2 knock-in line^47^, with *Clk*>CD8-GFP outlining clock-neuron membranes (**Figure 3G**). Me31B formed discrete cytoplasmic puncta in the large ventral lateral neurons (lLNv) at all four time points sampled. Three-dimensional segmentation showed that both the number (∼150 foci per lLNv) and total intensity of Me31B puncta were stable across ZT0, ZT6, ZT12, and ZT18 (**Figures 3H–3J**). P-bodies are therefore a constitutive feature of clock-neuron cytoplasm, present at constant abundance across the day.

We then performed three-color imaging of endogenous PER-mNG, Me31B-Tom2, and a *Clk*>BFP-NLS nuclear marker at ZT18, near the peak of cytoplasmic PER (**Figure 3K**). PER formed discrete cytoplasmic puncta, many in close apposition to ME31B-positive P-bodies: ∼50– 55% of PER foci in the lLNv were scored as adjacent (**Figure 3L**, see Methods). The PER and ME31B signals occupied distinct, non-overlapping puncta, indicating that PER is proximal to P-bodies rather than residing within these granules. This arrangement places PER close enough to ME31B-positive granules to permit proximity biotinylation, confirming by imaging the spatial relationship inferred from proximity labeling. Because P-bodies are canonical sites of mRNA storage and translational repression, we next asked whether the clock-gene mRNAs themselves, *per* and *tim*, localize to P-bodies in lLNv neurons.

### *per* and *tim* mRNAs localize to Me31B-labeled P-bodies, most prominently at ZT12

We followed *per* mRNA across the circadian cycle by HCR fluorescence *in situ* hybridization with probes against *per* exons, in brains co-labeled with *Clk*>CD8-GFP for clock-neuron membranes (**Figure 4A**). *per* mRNA abundance oscillated across the day, rising from ∼30 foci per lLNv at ZT0–8 to a peak of ∼115 foci at ZT16 (**Figure 4B**). To test whether *per* mRNA puncta associate with P-bodies, we performed three-color imaging of *per* mRNA, the P-body marker Me31B-GFP, and the lLNv marker PDF at multiple ZTs during the mRNA accumulation phase (**Figure 4C**). at ZT12, ∼55% of *per* mRNA puncta were associated with ME31B-positive foci (see Methods), falling to ∼30% at ZT16 and ∼32% at ZT20 (**Figure 4D**). Three-dimensional rendering of lLNVs cytoplasm showed the same pattern, with the number of *per* mRNAs within 0.4 µm of the nearest ME31B focus decreasing progressively from ZT12 to ZT20 (**Figure 4E**).

**Fig 4.**
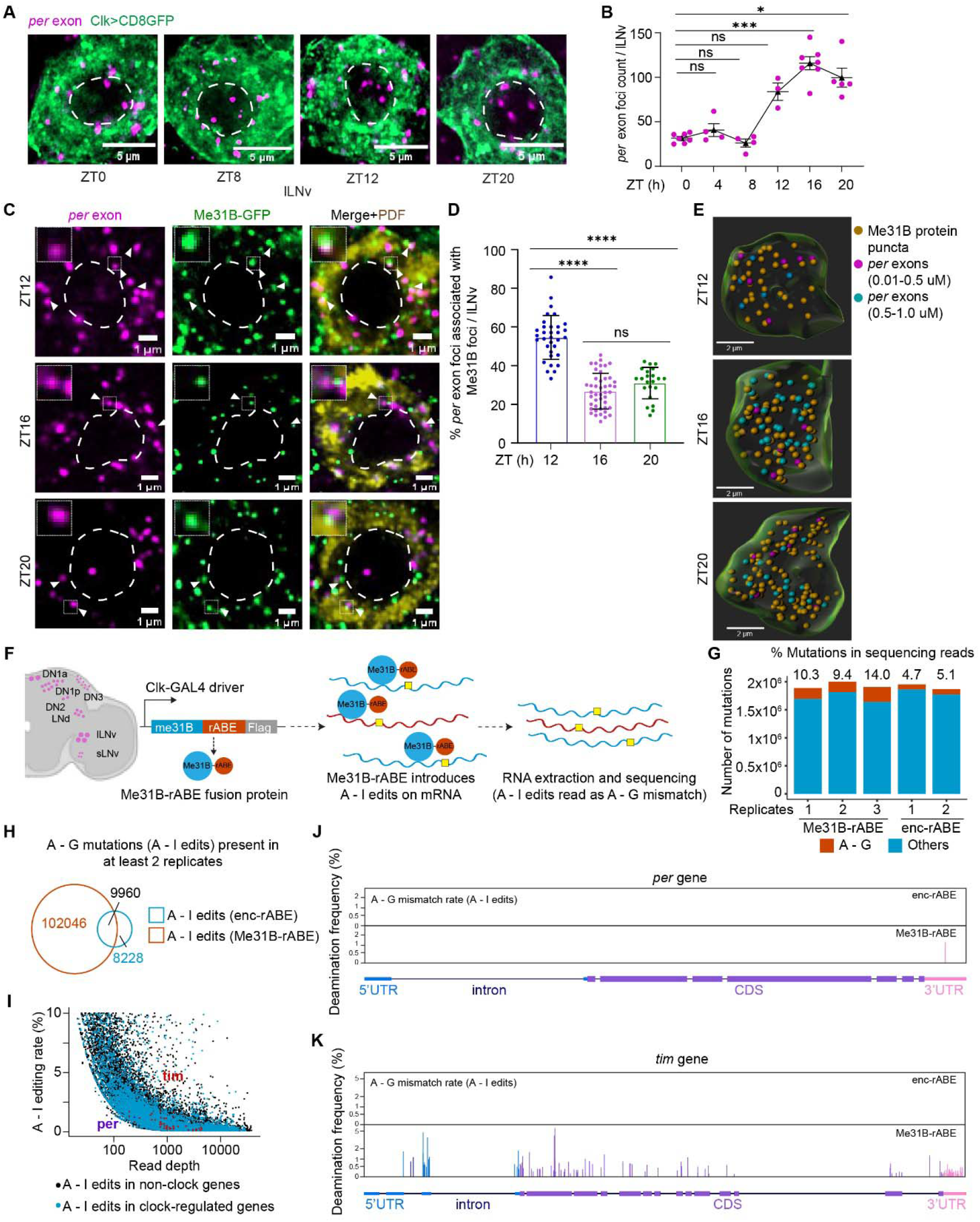
*per* and *tim* mRNAs associate with Me31B-positive P-bodies, most prominently at ZT12. **(A)** Representative images of *per* exonic HCR-FISH foci (magenta) in large ventral lateral neurons (lLNv) of *Clk*>CD8-GFP flies (green, membrane marker) at ZT0, ZT8, ZT12 and ZT20. Dashed lines demarcate nuclei. Scale bars, 5 µm. **(B)** Quantification of *per* foci per lLNv at ZT0, ZT4, ZT8, ZT12, ZT16 and ZT20. **(C)** Representative images of *per* exonic foci (magenta), Me31B-GFP (green) and PDF (yellow) in lLNv of Me31B-GFP knock-in flies at ZT12, ZT16 and ZT20. Arrowheads mark *per* foci associated with Me31B foci; insets show magnified examples. Dashed lines demarcate nuclei. Scale bars, 1 µm. **(D)** Percentage of *per* foci associated with Me31B foci at ZT12 (n = 34), ZT16 (n = 45) and ZT20 (n = 22). A focus was scored as associated when its centroid lay within 400 nm of the nearest Me31B focus. **(E)** Representative 3D renderings of lLNv at ZT12, ZT16 and ZT20, showing segmented Me31B puncta (brown) and *per* foci within 0.01–0.5 µm (magenta) or 0.5–1.0 µm (teal) of the nearest Me31B punctum. Scale bars, 2 µm. **(F)** Schematic of A-to-I editing by the Me31B-rABE fusion in clock neurons of *Clk*-GAL4 > *me31B*-rABE-FLAG flies, followed by cell sorting, RNA extraction and sequencing. **(G)** Percentage of A-to-G mismatches among total mismatches in Me31B-rABE (n = 3) and Enc-rABE control (n = 2) samples. **(H)** Venn diagram of A-to-I editing sites recovered in at least two replicates, showing Me31B-rABE-specific, Enc-rABE-specific and shared sites. **(I)** A-to-I editing rate versus read depth for clock-regulated (blue) and non-clock (black) genes; *per* and *tim* are labeled. **(J, K)** Gene-track views of A-to-I editing along *per* (J) and *tim* (K) in Enc-rABE and Me31B-rABE samples. Each data point represents a single neuron. lLNv, large ventral lateral neuron; ZT, Zeitgeber time; ns, not significant. Statistical tests and exact p-values are given in Table S3.

A complementary dataset using a Me31B-Tom2 knock-in line for *per* mRNA yielded the same pattern at ZT12 versus ZT18 (**Ext. Data Figures 4A** and **4B**), and *tim* mRNA also associated with P-bodies (∼35% at ZT12; **Ext. Data Figures 4C and 4D**). Across these time points, the associated fraction fell while total *per* transcript rose: the P-body-associated pool changed little in absolute terms (∼47, ∼35, and ∼34 foci at ZT12, ZT16, and ZT20) even as total *per* increased from ∼85 to ∼115 foci. The associated fraction was highest at ZT12, on the rising arm of *per* accumulation and well before peak nuclear PER (ZT18–24), consistent with P-bodies harboring a pool of clock-gene transcripts during early RNA accumulation.

### *per* and *tim* transcripts are edited by Me31B-rABE in clock neurons

Next, to molecularly footprint Me31B–RNA associations in clock neurons, we applied the rABE proximity-editing approach^48^, in which an engineered RNA adenosine deaminase fused to an RNA-binding protein deposits adenosine to inosine (A-to-I) edits on transcripts within molecular contact distance of the fusion partner, providing a permanent record read out as A-to-G mismatches in standard RNA sequencing. We generated UAS-Me31B-rABE-Flag transgenic flies and drove expression with *Clk*-GAL4, restricting editing to clock neurons of the adult brain (**Figure 4F**). As a specificity control, we generated UAS-Enc-rABE-Flag flies, in which the same editor is fused to Encore, an RNA-binding protein that does not localize to P-bodies, and expressed it with the same driver. Clock neurons were isolated by fluorescence-activated cell sorting, and RNA from three biological replicates of Me31B-rABE flies, and two replicates of Enc-rABE flies was sequenced. Because editing is confined to the sorted population, A-to-G signal reports ME31B contacts within clock neurons directly rather than against a background of unedited transcripts from other cell types. The A-to-G fraction of total read mismatches was elevated to 9.4–14.0% in Me31B-rABE samples versus 4.7–5.1% in controls (**Figure 4G**), indicating rABE editing activity above the endogenous ADAR background.

To call high-confidence editing sites, we adapted the published analysis pipeline^48^: filtering for coverage (≥20 average reads per site), removing positions with detectable A-to-G signal in Enc-rABE controls (allele count ≥2) to eliminate genomic SNPs and endogenous A-to-I edits, and applying the Cochran–Mantel–Haenszel test on per-position allele-count contingency tables across replicates with Benjamini–Hochberg correction (p_adj < 0.05). Sites recovered in at least two Me31B-rABE replicates were retained. The pipeline returned 102,046 Me31B-rABE-specific sites, versus 8,228 control-only and 9,960 shared sites (**Figure 4H**).

Editing rates spanned ∼0.1% to >50% (**Figure 4I and Ext. Data Figure 4E**), with the high-rate tail dominated by sites also present in control samples, consistent with endogenous ADAR activity. Me31B-rABE-specific sites were concentrated in CDS and 3′UTR regions (**Ext. Data Figure 4F**), consistent with cytoplasmic mRNA targeting and with the reported binding distribution of the mammalian ortholog DDX6^48^. The abundant transcript Act5C, a previously reported Me31B target^48^, was edited in Me31B-rABE but not Enc-rABE samples (**Ext. Data Figure 4G**), confirming that the fusion recovers known Me31B-associated mRNAs. Clock-regulated transcripts were edited across the same range as the transcriptome as a whole (**Figure 4I**, blue), indicating that Me31B contacts the clock-neuron transcriptome broadly rather than only clock genes. At single-gene resolution, *tim* showed extensive editing across its CDS and UTRs (**Figure 4K**), whereas editing of *per* was more restricted, concentrated at a discrete 3′UTR position (**Figure 4J**).

Per-site editing rates were low for most transcripts, including *per* and *tim*, once the endogenous ADAR-derived high-rate tail is excluded. This is expected, because deamination is stochastic: each Me31B–RNA encounter has a low probability of editing any individual adenosine, so a transcript may be contacted repeatedly yet edited at only a subset of positions. Together with the imaging data above, these results indicate that per and tim transcripts associate with Me31B in clock neurons.

### Ribosome profiling reveals low translational efficiency of *per* and *tim* at ZT12

Because P-bodies are canonical sites of translational repression, we asked whether *per* and *tim* transcripts are poorly translated at ZT12, the phase of maximal P-body association, by performing clock-neuron-specific ribosome profiling. We adapted a tissue-specific ribosome profiling strategy^49^: UAS-RpL3-FLAG was driven by *tim*-GAL4; intact ribosomes were affinity-purified from adult brain lysate with anti-FLAG magnetic beads; ribosome-protected mRNA fragments were liberated by RNase digestion, size-selected, and sequenced; and total RNA from the same heads was sequenced in parallel as a transcriptional reference (three biological replicates, **Figure 5A**). Because ribosome-protected fragments are recovered only from cells expressing the tagged subunit, this approach yields ribosome-occupancy data restricted to clock neurons even when using whole-brain input. Translational efficiency (TE) for each gene was computed as the ratio of ribosome-protected to total RNA reads.

**Fig. 5.**
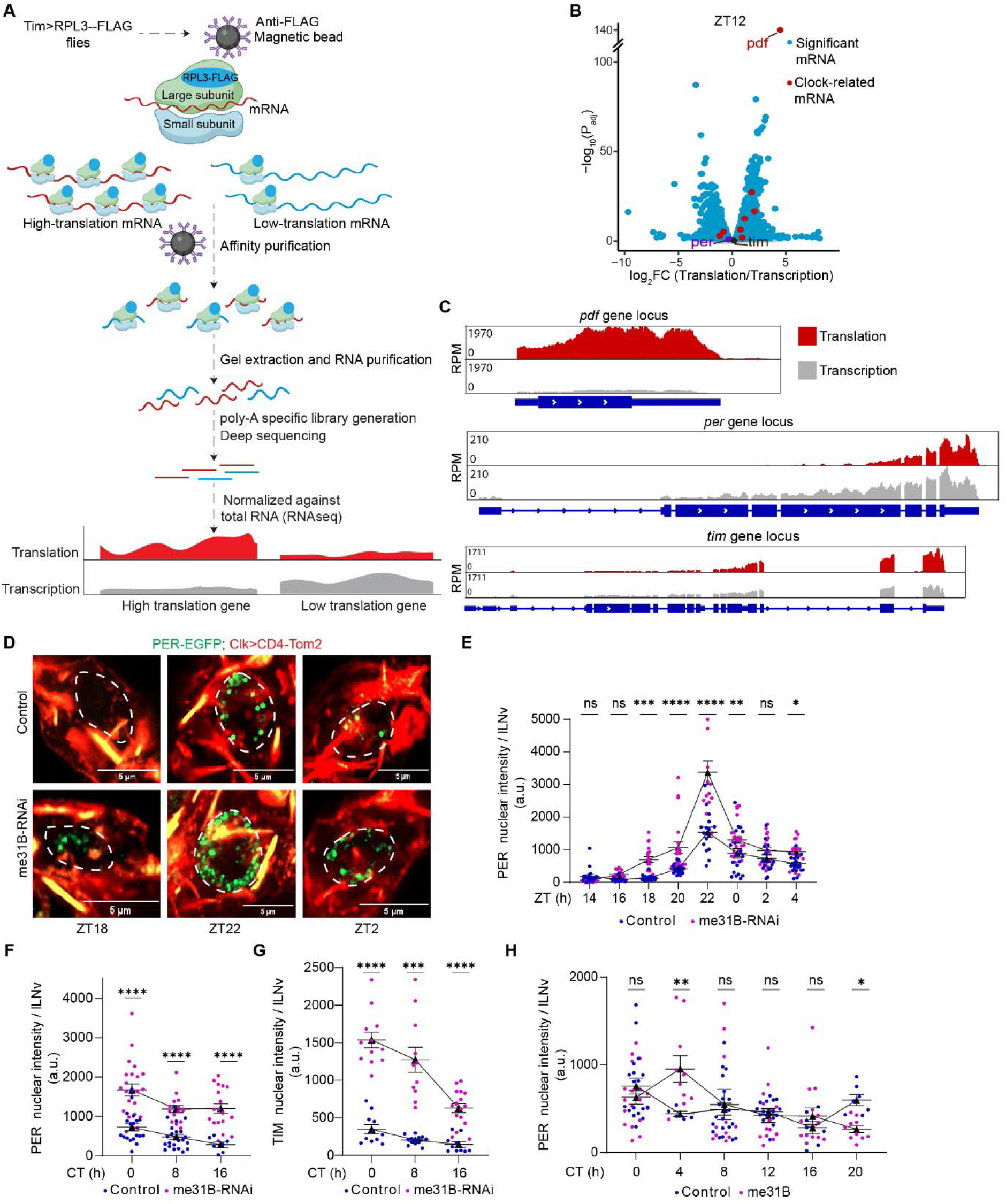
*per* and *tim* show low translational efficiency at ZT12, and Me31B levels set the timing and amplitude of PER accumulation. (A) Schematic of the ribosome profiling strategy. FLAG-tagged RpL3 was expressed in clock neurons of *tim*-GAL4>UAS-RpL3-FLAG flies and ribosomes were affinity-purified from head lysates with anti-FLAG beads. Ribosome-protected fragments were size-selected and sequenced, and read counts were normalized to total RNA from the same lysates to give translational efficiency. (B) Volcano plot of the clock-neuron translatome at ZT12, plotted as log (translation/transcription) against −log (P_adj). Blue, significantly enriched or depleted transcripts; red, clock-related transcripts. *pdf*, *per* and *tim* are labeled. (C) Gene-track views of ribosome-protected fragment reads (translation, red) and total RNA reads (transcription, grey) at the *pdf* (top), *per* (middle) and *tim* (bottom) loci. RPM, reads per million mapped reads. (D) Representative images of PER-EGFP foci (green) in large ventral lateral neurons (lLNv) of *per*-EGFP; *Clk*>CD4-tdTomato (control, top) and *per*-EGFP; *Clk*>CD4-tdTomato; *me31B*-RNAi (bottom) flies at ZT18, ZT22 and ZT2. Membranes are marked by CD4-tdTomato (red); dashed lines demarcate nuclei. Scale bars, 5 µm. (E) PER-EGFP nuclear intensity per lLNv in control and *me31B*-RNAi flies across ZT14–ZT4, quantified from (D). (F) PER-EGFP nuclear intensity per lLNv in control and *me31B*-RNAi flies at CT0, CT8 and CT16 in constant darkness. (G) TIM-mNeonGreen nuclear intensity per lLNv in *tim*-mNeonGreen; *Clk*>CD4-tdTomato (control) and *tim*-mNeonGreen; *Clk*>CD4-tdTomato; *me31B*-RNAi flies at CT0, CT8 and CT16. (H) PER-EGFP nuclear intensity per lLNv in control and *Clk*>UAS-Me31B (Me31B overexpression) flies across CT0–CT20. Each data point represents a single neuron. lLNv, large ventral lateral neuron; ZT, Zeitgeber time; CT, circadian time; FC, fold change; ns, not significant; a.u., arbitrary units. Statistical tests and exact p-values are given in Table S3.

At ZT12, the clock-neuron translatome contained thousands of mRNAs whose TE values spanned more than three orders of magnitude (**Figures 5B** and **Ext. Data Figures 5A** and **5B**). The neuropeptide *pdf*, constitutively and abundantly translated in lLNv neurons, served as an internal positive control and was recovered as the most strongly translation-enriched transcript in the dataset (**Figure 5B**; gene-track view in **Figure 5C**), whereas *ND1* showed low translation enrichment despite high transcript abundance (**Ext. Data Figure 5C**). By contrast, *per* and *tim* were abundantly transcribed at ZT12 yet showed no translational enrichment, with ribosome occupancy tracking rather than exceeding transcript abundance (log FC ≈ 0; **Figure 5B**, red points; gene tracks in **Figure 5C**). Abundant transcript with no corresponding translational enrichment is the pattern expected if these mRNAs are withheld from translation, consistent with their association with Me31B-labeled P-bodies at this phase.

### Bidirectional perturbation of *me31B* alters PER accumulation timing and the circadian period

Next, we tested the effects of reducing and enhancing Me31B activity in clock neurons using *Clk*-GAL4-driven *me31B-RNAi* (loss of function) and *UAS-Me31B* (gain of function). *me31B*-RNAi reduced total Me31B-GFP intensity in lLNv neurons by ∼50% and eliminated discrete ME31B puncta, leaving only diffuse cytoplasmic signal (**Ext. Data Figures 5D and 5E**). Loss of P-bodies upon ME31B depletion is consistent with the requirement for DDX6 in P-body assembly in mammalian cells^43^, and indicates that the knockdown removes the compartment itself rather than only reducing ME31B levels. As shown above, *me31B* knockdown rendered most flies arrhythmic (8.5% rhythmic; **Figure 3C**). Co-expressing UAS-Me31B with *me31B*-RNAi rescued the behavioral phenotype, restoring rhythmicity to 83.3% and a normal period of 23.63 ± 1.29 h (**Ext. Data Figure 5F**), indicating that the behavioral defects are caused by loss of Me31B.

In control flies maintained in 12:12 light:dark (LD) cycles, PER-EGFP nuclear intensity in lLNv neurons rose from baseline at ZT18 to a peak at ZT22 and declined by ZT4, reproducing the canonical late-night peak of nuclear PER (**Figure 5D**, top row; **Figure 5E**, blue trace). *me31B-RNAi* flies showed two distinct deviations from this pattern: PER appeared prematurely in the nucleus at ZT18 (**Figure 5D**, bottom row; **Figure 5E**, magenta trace), and the eventual ZT22 peak overshot control levels by approximately 2-fold, indicating that Me31B constrains both the onset and the magnitude of PER accumulation. In constant darkness (DD), *me31B*-RNAi flies showed elevated nuclear PER and TIM at all sampled timepoints relative to controls (**Figures 5F and 5G**, magenta versus blue traces; **Ext. Data Figure 5G**).

The reciprocal manipulation produced the opposite phenotype. In *Clk*>*UAS-Me31B* flies, free-running locomotor rhythms were lengthened to a period of 26.55 ± 0.19 h (78.57% rhythmicity; **Ext. Data Figure 5H**), representing a ∼2-h period lengthening relative to control. Consistent with this behavioral phenotype, PER nuclear accumulation on the second day of constant darkness (DD2) was delayed by ∼2 h across the circadian cycle relative to control (**Figure 5H**). Together, these bidirectional perturbations indicate that Me31B levels set the timing and amplitude of PER accumulation: loss of Me31B causes PER to appear early and accumulate to higher levels, and abolishes behavioral rhythmicity, whereas excess Me31B delays PER accumulation and lengthens the circadian period.

### Pcm depletion elevates nuclear PER and TIM and extends the repression phase

In our RNAi screen of PER-proximal candidates (**Figure 3**), the 5′–3′ exonuclease Pcm, which mediates bulk cytoplasmic mRNA decay, emerged as a strong hit (**Figure 3F**). We therefore asked whether *pcm* knockdown alters clock-protein accumulation across the circadian cycle, driving UAS-*pcm*-RNAi with *Clk*-GAL4 and quantifying PER, TIM and CLK in DD. PER-EGFP nuclear intensity in control lLNv neurons cycled across the day, high at CT0 and declining to a trough at CT16–CT20 (**Figure 6A**, top row; **Figure 6B**, blue trace). *pcm-RNAi* flies showed a markedly different pattern: PER nuclear intensity remained elevated at every phase sampled (**Figure 6A**, bottom row; **Figure 6B**, magenta trace). The phenotype was more striking still when we counted discrete PER nuclear foci. In control flies, PER foci cycled with a peak of ∼14 foci/lLNv at CT4 and decreased to ∼2 foci by CT12–CT16 (**Figure 6C**, blue trace). In *pcm-RNAi*, foci accumulated to ∼22 foci/lLNv at CT8 and remained elevated at CT12 and CT16, phases at which control foci have largely disassembled (**Figure 6C**, magenta trace). TIM behaved similarly: TIM-mNeonGreen nuclear intensity declined across the subjective day in controls (**Figure 6D**, top row; **Figure 6E**, blue trace), whereas *pcm-RNAi* flies maintained two- to four-fold elevated TIM at every phase (**Figure 6D**, bottom row; **Figure 6E**, magenta trace).

**Fig. 6.**
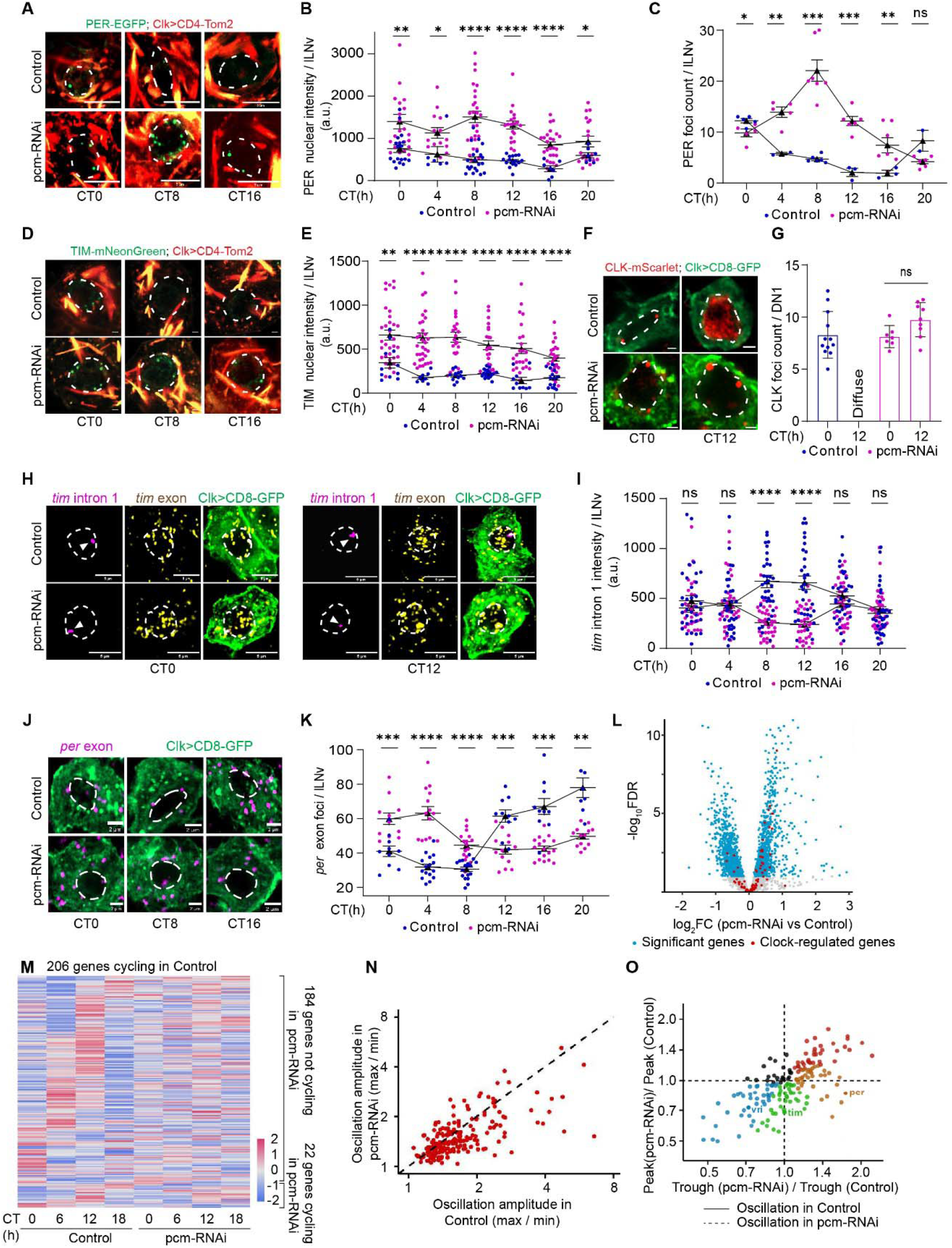
Knockdown of *pcm* prolongs nuclear PER and TIM accumulation and extends the repression phase. **(A)** Representative images of PER-EGFP foci (green) in large ventral lateral neurons (lLNv) of *per*-EGFP; *Clk*>CD4-tdTomato (control, top) and *per*-EGFP; *Clk*>CD4-tdTomato; *pcm*-RNAi (bottom) flies at CT0, CT8 and CT16. Membranes are marked by CD4-tdTomato (red); dashed lines demarcate nuclei. Scale bars, 5 µm. **(B, C)** PER-EGFP nuclear intensity (B) and nuclear foci count (C) per lLNv across CT0–CT20, quantified from (A). **(D)** Representative images of TIM-mNeonGreen (green) in lLNv of *tim*-mNeonGreen; *Clk*>CD4-tdTomato (control) and *tim*-mNeonGreen; *Clk*>CD4-tdTomato; *pcm*-RNAi flies at CT0, CT8 and CT16. Scale bars, 5 µm. **(E)** TIM-mNeonGreen nuclear intensity per lLNv across CT0– CT20, quantified from (D). **(F)** Representative images of CLK-mScarlet (red) in DN1 neurons of *clk*-mScarlet; *Clk*>CD8-GFP (control) and *clk*-mScarlet; *Clk*>CD8-GFP; *pcm*-RNAi flies at CT0 and CT12. Scale bars, 5 µm. **(G)** CLK-mScarlet nuclear foci count per DN1 at CT0 and CT12, quantified from (F). In controls at CT12, CLK signal was diffuse rather than punctate. **(H)** Representative images of *tim* intron 1 (magenta) and *tim* exonic (yellow) HCR-FISH signal in lLNv of *Clk*>CD8-GFP (control) and *Clk*>CD8-GFP; *pcm*-RNAi flies at CT0 and CT12. Scale bars, 5 µm. **(I)** *tim* intron 1 intensity per lLNv across CT0–CT20, quantified from (H). **(J)** Representative images of *per* exonic HCR-FISH foci (magenta) in lLNv of *Clk*>CD8-GFP (control) and *Clk*>CD8-GFP; *pcm*-RNAi flies at CT0, CT8 and CT16. Scale bars, 2 µm. **(K)** *per* foci per lLNv across CT0–CT20, quantified from (J). **(L)** Volcano plot of differential gene expression between control and *pcm*-RNAi FACS-purified clock neurons. Blue, significantly changed genes; red, clock-regulated genes. **(M)** Heat map of the 206 genes scored as cycling in controls, showing expression across CT0, CT6, CT12 and CT18 in control and *pcm*-RNAi samples. Of these, 184 lost detectable rhythmicity in *pcm*-RNAi and 22 continued to cycle. **(N)** Oscillation amplitude (max/min) of cycling genes in *pcm*-RNAi versus control. Dashed line, unity. **(O)** Peak and trough ratios (*pcm*-RNAi/control) for cycling genes; *per*, *tim* and *vri* are labeled. Dashed lines mark unity on each axis. Each data point in (B, C, E, G, I, K) represents a single neuron. lLNv, large ventral lateral neuron; DN1, dorsal neuron 1; CT, circadian time; FC, fold change; ns, not significant; a.u., arbitrary units. Statistical tests and exact p-values are given in Table S3.

**Fig. 7.**
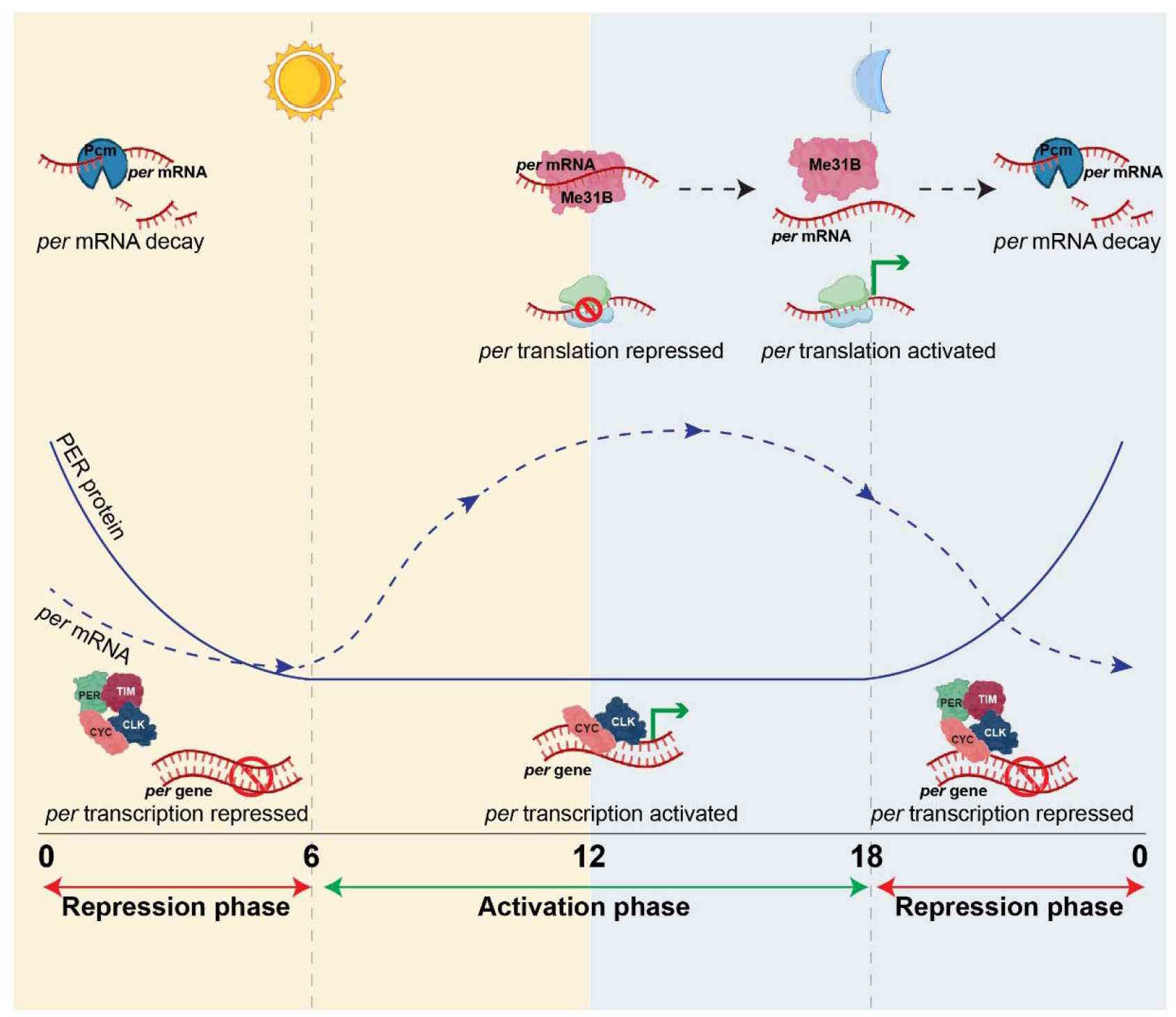
Model for P-body-mediated control of *per* mRNA across the circadian cycle. During the activation phase, CLK/CYC drives *per* transcription and newly made transcripts accumulate in Me31B-labeled P-bodies, where they are held in a translationally repressed state. Release from Me31B permits translation, so PER protein accumulates several hours after its transcript, generating the delay required for sustained oscillation. During the repression phase, PER and TIM enter the nucleus to shut down CLK/CYC-dependent transcription, and the 5′–3′ exonuclease Pcm degrades remaining *per* transcripts, clearing the pool so that repression ends on schedule. Solid line, PER protein; dashed line, *per* mRNA.

The persistence of nuclear PER and TIM, together with the failure of PER foci to disassemble, suggested that Pcm depletion extends the repression state. We tested this at the level of the activator complex by imaging CLK-mScarlet in DN1 neurons, where CLK-mScarlet signal is brightest and where the CLK foci cycle has been previously characterized^9^. In controls, CLK was organized into ∼8 discrete nuclear foci at CT0 and became diffusely distributed by CT12 (**Figure 6F**, top row; **Figure 6G**), as described previously^9^. In *pcm-RNAi* flies, CLK remained in discrete foci at CT12 (∼10 foci at CT12 vs ∼8 foci at CT0; **Figure 6F**, bottom row; **Figure 6G**), a configuration normally restricted to the repression phase. Together, elevated nuclear PER and TIM, persistent PER foci, and retention of CLK in foci indicate that Pcm depletion extends the repression phase.

### Pcm depletion suppresses nascent transcription and dampens clock-controlled gene cycling

If *pcm*-RNAi extends the repression state, nascent transcription of CLK/CYC target genes should be suppressed throughout most of the day. To assay nascent transcription rather than steady-state mRNA, we performed HCR-FISH against the first intron of *tim*, which is rapidly degraded following co-transcriptional splicing and therefore reports active transcription at the time of fixation^50, 51^ (**Figure 6H**). In control lLNv neurons, *tim* intron 1 signal cycled with a peak at CT8–CT12, reflecting the canonical CLK/CYC-driven activation phase (**Figure 6I**, blue trace). In *pcm-RNAi* flies, signal at CT8 and CT12 fell to less than half of control values and remained low throughout the cycle, with only a modest residual rise toward the end of the day (**Figure 6I**, magenta trace). This decrease in nascent transcription at the canonical activation phase is the expected consequence of a prolonged repression phase.

Impaired decay combined with suppressed transcription predicts a characteristic distortion of steady-state clock-gene mRNA: reduced peaks, because new synthesis is suppressed, and elevated troughs, because existing transcripts fail to clear. To test this, we performed HCR-FISH against *per* exons in lLNv neurons (**Figure 6J**). In controls, *per* mRNA cycled from a trough of ∼30 foci/lLNv at CT8 to a peak of ∼80 foci at CT20 (**Figure 6K**, blue trace). In *pcm-RNAi*, both peak and trough were altered and phase-shifted: the trough rose to ∼45 foci (at CT12, versus ∼30 at CT8 in controls), and the peak fell to ∼62 foci (at CT4, versus ∼80 at CT20 in controls), with the *per* trajectory differing from control at every phase sampled (**Figure 6K**, magenta trace).

To ask whether these changes generalize across the clock-controlled transcriptome, we performed bulk RNA-seq using fly heads at CT0, CT6, CT12, and CT18 in control and *pcm-RNAi* flies (3 biological replicates per condition). Differential-expression analysis identified many transcripts, including clock-regulated genes, with altered abundance between conditions (**Figure 6L**). Of 206 genes scored as cycling in controls, 184 (89%) lost detectable rhythmicity in *pcm*-RNAi (**Figure 6M**), and oscillation amplitudes were dampened among those that remained (**Figure 6N**). Decomposing the amplitude loss into peak and trough components revealed that rhythms are suppressed through multiple distinct modes (**Figure 6O**). Key clock transcripts, including *per* and *tim*, fell in the elevated-trough, reduced-peak quadrant (**Figure 6O**, lower right), consistent with the imaging results. Loss of Pcm therefore collapses the clock-controlled transcriptome, through both loss of rhythmicity and peak–trough distortion of the transcripts that continue to cycle.

## Discussion

Every transcription–translation feedback oscillator must separate the accumulation of a repressor’s transcript from the action of its protein, yet in the circadian clock this lag has been explained mostly through the post-translational regulation of the repressor protein. Our results identify a step that acts earlier, on the transcript itself: *per* and *tim* are held in Me31B-labeled P-bodies as they accumulate, where they are poorly translated; Me31B levels set when and how much PER accumulates; and Pcm is required for repression to end on time. To date, the known cytoplasmic regulators operate after synthesis: DOUBLETIME and CASEIN KINASE 2 phosphorylate PER to control its stability and the timing of its nuclear entry^14, 15^, and SLIMB-mediated ubiquitination sets its turnover^16–18^. *per* and *tim* are abundantly transcribed at ZT12 yet show low translational efficiency at precisely the phase when their association with Me31B-positive P-bodies is highest, the signature expected of transcripts held in a translationally silent state. We propose that this mechanism acts in series with phosphorylation-dependent control rather than as an alternative to it: sequestration influences when repressor synthesis begins, whereas phosphorylation determines how long the resulting protein persists and when it enters the nucleus. Consistent with a step that acts upstream, P-body association peaks hours before nuclear PER does.

Intriguingly, PER foci are adjacent to, but do not overlap with, Me31B-positive granules, placing the repressor protein at the surface of the compartment that holds its own transcript. Whether PER contributes to regulating that compartment, or is simply recruited alongside its own mRNA, is not resolved by our data.

### Capture and clearance act on the same problem from opposite ends

Two features of the P-body compartment explain how it can impose a delay. Me31B granules are constitutive, stable in number (∼150 per lLNv) and intensity across the day, and the P-body-associated per pool changes little in absolute terms (∼47, ∼35 and ∼34 foci at ZT12, ZT16 and ZT20) while total per rises steeply. Sequestration is therefore not proportional to transcript abundance: the fraction withheld from translation is greatest early in accumulation, when withholding matters most for setting the delay. Bidirectional genetic tuning shows that Me31B levels are limiting for repressor timing. Knockdown causes premature PER accumulation that overshoots control levels ∼2-fold and abolishes rhythmicity, whereas overexpression delays nuclear PER by ∼2 h and lengthens the free-running period to 26.6 h while preserving robust rhythms; the knockdown phenotype is rescued by Me31B co-expression. A technical limitation of this perturbation is that *me31B* knockdown simultaneously eliminates physical P-bodies and removes Me31B’s enzymatic activity, preventing us from fully disentangling the contribution of the physical compartment from that of the helicase itself. Loss of Pcm produces the complementary defect at the other end of the transcript life cycle: repressor proteins persist, the repression phase is extended, nascent *tim* transcription is reduced, and 89% of cycling transcripts in clock neurons lose rhythmicity. Because each factor was perturbed independently, we cannot establish a physical or temporal link between them; whether Pcm acts on the transcript pool held by Me31B or on a distinct cytoplasmic pool remains an open question.

### A repressive arm complementing translational activation

Past studies of cytoplasmic clock RNA regulation in clock neurons have largely focused on translational activators such as ATX2, TYF, and PABP, which promote *per* translation^19–21^. Our results place ME31B in a repressive relationship with the core loop transcripts themselves, which differs from a previous report that the ATX2–NOT1 complex containing ME31B does not affect PER translation and acts through a PER-independent route on clock outputs^26^. Two factors could reconcile these findings. First, our measurements are time-resolved and report the phase of PER accumulation, a shift that a fixed-timepoint or reporter-based assay would not detect. Second, ME31B’s activity is dictated largely by its binding partners^42–44^. While it associates with ATX2 and NOT1 in certain contexts, it associates with EDC3, PATR-1, LSM3 and GE-1, all recovered here as PER-proximal P-body components, in other contexts. A timed exchange of cofactors, rather than a shift in helicase abundance, could serve as the switch that releases the translational brake, making brake release itself a regulated step. What confers transcript selectivity remains to be determined.

Proximity labeling reports molecular neighborhoods within ∼10 nm, so enrichment does not distinguish direct interactors from proteins sharing a compartment. We therefore present the 252-protein proximitome as a spatial neighborhood resource rather than a direct interaction map.. DDX6 and XRN1 are among the most broadly conserved eukaryotic RNA-binding proteins, and the P-body functions they carry out, translational storage and 5′–3′ decay, are shared from yeast to mammals. Ribosome profiling has shown that the translational efficiency of many transcripts oscillates diurnally, largely independently of mRNA abundance, in organisms ranging from *Neurospora* to mammals^52–54^. Sequestration of transcripts in an RNP compartment provides a potential mechanism for this uncoupling. Whether DDX6-dependent sequestration times *Per* and *Cry* translation in the suprachiasmatic nucleus remains to be tested. More broadly, a constitutive RNP compartment that holds back a rising transcript, paired with a decay enzyme that clears it, offers a general way to build a multi-hour delay into biological negative-feedback loops.

## Materials and Methods

### Fly Strains

All Drosophila melanogaster strains were maintained on standard cornmeal-yeast-agar medium at 25°C under a 12-hour light/dark cycle. The fly strains used in the study were previously described or obtained from the Bloomington Stock Center: *w*^1118^ (BL-6326), *y,sc,v* (BL-25709), *Clk-GAL4*^55^, *tim*-UAS-GAL4 (TUG) (BL-80941), *UAS-Dcr2* (BL-24644), *UAS-CD8GFP*^56^, *UAS-CD4tdTom*^56^, *per-mNeonGreen*^9^, *tim-mNeonGreen*^57^, *per-EGFP*^9^, *clk-mScarlet*-I^9^, *Me31B-Tom2*^47^, *Me31B-GFP*^47^, *UAS-me31B*^58^, *UAS-RpL3-FLAG*^49^. The following lines were generated in this study: *per*-V5-miniTurbo, *UAS-Me31B-rABE-Flag*. All the RNAi lines used in the study are described in Table S2. The locomotor activity rhythms data and genotype descriptions are presented in Table S2. For control experiments, GAL4 and UAS lines are crossed to w1118 flies (the genetic background of GAL4 and RNAi lines). For behavioral experiments, males were used, except for the *per ¹*/*per*-V5-mTb rescue, in which females were used. For imaging and sequencing experiments, both males and females were used

### CRISPR protocol for generating *per*-V5-miniTurbo flies

The *per*-V5-miniTurbo knock-in was generated using the same two sgRNAs and the same 5′ and 3′ homology arms described previosuly for *per*-mNeonGreen^9^, with the mNeonGreen cassette replaced by a V5-miniTurbo cassette. The cassette, comprising a V5 epitope tag and the miniTurbo coding sequence (based on Addgene Plasmid 107168), was designed to be fused in frame to the C-terminus of PER with the native *per* stop codon removed. Donor plasmid assembly, transformation, colony screening and sequence verification were performed as described for *per*-mNeonGreen. Rainbow Transgenic Flies, Inc. performed injections into *nos*-Cas9 attP2 embryos.

Injected adults were crossed individually to FM7a males or virgin females. F1 progeny were screened by PCR for the insert using the leg-lysis protocol described previously^9, 59^. Flies carrying the insertion were backcrossed to FM7a and their progeny homozygosed. Homozygous stocks were genotyped and Sanger-sequenced.

### Transgenic constructs

The following fly lines were created for this study: UAS-Me31B-rABE-FLAG and UAS-Enc-rABE-FLAG. To identify transcripts associated with ME31B in clock neurons, we generated an overexpression line encoding a C-terminal fusion of the rABE editor and a FLAG tag to the Me31B coding sequence (UAS-Me31B-rABE-FLAG). UAS-Enc-rABE-FLAG served as the specificity control, since Encore is an RNA-binding protein that does not localize to P-bodies. In both constructs, the coding sequence was inserted in frame N-terminal to the rABE editor followed by a 3×FLAG tag, separated by a linker sequence. Constructs were cloned into the pJFRC-MUH expression plasmid (Addgene #26213) in the default 10×UAS configuration. Plasmids were cloned from gBlock fragments (Integrated DNA Technologies) using Gibson assembly (NEBuilder HiFi DNA Assembly Master Mix, New England Biolabs) following the manufacturer’s protocol. Clones were verified by whole-plasmid sequencing (Eurofins), prepared for injection by midiprep (QIAGEN) and microinjected into a phiC31 integrase line carrying the attP40 docking site (Rainbow Transgenic Flies). Transformants were crossed individually to a balancer line, backcrossed and homozygosed.

### Protein proximity labeling

#### Sample collection

Flies were maintained on standard cornmeal-yeast-agar food without prior biotin depletion. To label proteins in the vicinity of PER-V5-miniTurbo, 7-day-old adults entrained to a 12:12 light:dark cycle at 25 °C were transferred to food supplemented with 100 µM D-biotin (Sigma-Aldrich B4504) dissolved directly into the standard fly food mixture. Flies were fed on biotin-supplemented food for a single 6-h window, timed to one of four Zeitgeber phases (ZT0–6, ZT6–12, ZT12–18 or ZT18–24), and collected immediately at the end of the window. Control flies carrying the per-mNeonGreen knock-in, which lacks biotin ligase activity, were fed identical biotin-supplemented food over the ZT0–6 window to account for endogenously biotinylated proteins and non-specific bead binding. After feeding, flies were anesthetized on a CO pad, transferred to a 15 mL conical tube and flash-frozen in liquid nitrogen. Frozen flies were vortexed to separate heads from bodies at the brittle neck joint, and the mixture was passed through a mesh sieve to isolate heads, which were collected into microcentrifuge tubes and stored at −80 °C until processing. Approximately 300 heads were collected per sample.

#### Sample lysis

Frozen heads were homogenized in 1 mL ice-cold RIPA lysis buffer (Pierce 89900) supplemented with cOmplete protease inhibitor cocktail, EDTA-free (Roche 11873580001), using a motorized pellet pestle (Fisher Scientific K749540-0000 and 12-141-364). Homogenates were sonicated on ice (3 cycles of 10 s on/10 s off at 15% power; Fisher Scientific Sonic Dismembrator Model 500) and cleared by centrifugation at 16,000 × g for 15 min at 4 °C. Supernatants were transferred to fresh tubes.

#### Streptavidin pull-down

Cleared lysates were incubated with pre-equilibrated streptavidin-coated magnetic beads (Pierce Streptavidin Magnetic Beads, Thermo Fisher) overnight at 4 °C with gentle rotation. Beads were collected on a magnetic rack and washed sequentially at room temperature (twice, 5 min each) in 2% SDS in 50 mM Tris-HCl (pH 7.4), high-salt detergent wash buffer (0.1% sodium deoxycholate, 1% Triton X-100, 500 mM NaCl, 1 mM EDTA, 50 mM HEPES, pH 7.5), and lithium chloride wash buffer (250 mM LiCl, 0.5% NP-40, 0.5% sodium deoxycholate, 1 mM EDTA, 10 mM Tris-HCl, pH 8.0). For streptavidin blotting, bound proteins were eluted by boiling in 2× Laemmli sample buffer supplemented with 2 mM D-biotin and 20 mM DTT at 95 °C for 10–15 min. For LC-MS/MS analysis, beads were washed three additional times in 50 mM ammonium bicarbonate (pH 8.0) to remove residual detergents before undergoing on-bead trypsin digestion.

#### SDS-PAGE and streptavidin blotting

Protein samples were resolved on 4–20% gradient SDS-polyacrylamide gels (Bio-Rad Mini-PROTEAN TGX) at 200 V for 45 min in Tris-glycine-SDS running buffer, with a pre-stained protein ladder (Thermo Fisher PageRuler) for molecular-weight calibration. Proteins were transferred to PVDF membranes (Bio-Rad) using a wet transfer system (0.5 A, 25 min). Membranes were blocked in 5% BSA in TBST (0.1% Tween-20) for 1 h, then probed with streptavidin-HRP (1:5,000; Invitrogen) in blocking buffer for 1 h at room temperature. After three TBST washes (10 min each), blots were developed with ECL substrate (Pierce SuperSignal West Pico) and imaged on a ChemiDoc system (Bio-Rad).

#### Protein digestion and TMT labeling

Samples were proteolysed and labeled with TMT 10-plex according to the manufacturer’s protocol (Thermo Fisher). Briefly, cysteines were reduced (5 mM DTT, 30 min at 45 °C) and alkylated (15 mM 2-chloroacetamide, 30 min at room temperature), and proteins were precipitated by adding 6 volumes of ice-cold acetone followed by overnight incubation at −20 °C. The precipitate was pelleted by centrifugation and air-dried. The pellet was resuspended in 0.1 M TEAB and digested overnight (∼16 h) with trypsin/Lys-C mix (1:25 protease:protein; Promega) at 37 °C with constant mixing in a thermomixer. TMT 10-plex reagents were dissolved in 41 µL of anhydrous acetonitrile, and labeling was performed by transferring the entire digest to the TMT reagent vial and incubating at room temperature for 1 h. The reaction was quenched by adding 8 µL of 5% hydroxylamine and incubating for a further 15 min. Labeled samples were mixed and dried in a vacuum concentrator. The combined sample (∼200 µg) was fractionated into 8 fractions using a high-pH reversed-phase peptide fractionation kit according to the manufacturer’s protocol (Pierce, 84868). Fractions were dried and reconstituted in 9 µL of 0.1% formic acid/2% acetonitrile for LC-MS/MS analysis. TMT channel assignments are shown in Figure 1G.

#### Liquid chromatography–mass spectrometry analysis (LC-multinotch MS3)

To improve quantitation accuracy, we used multinotch-MS3^60^, which minimizes reporter-ion ratio distortion arising from fragmentation of co-isolated peptides. Data were acquired on an Orbitrap Fusion (Thermo Fisher Scientific) coupled to an RSLC Ultimate 3000 nano-UPLC (Dionex). Sample (2 µL) was resolved on a PepMap RSLC C18 column (75 µm i.d. × 50 cm; Thermo Fisher Scientific) at a flow rate of 300 nL/min using a 0.1% formic acid/acetonitrile gradient (2–22% acetonitrile over 150 min; 22–32% acetonitrile over 40 min; 20 min wash at 90%, followed by 50 min re-equilibration) and sprayed directly into the mass spectrometer using an EasySpray source (Thermo Fisher Scientific). The mass spectrometer collected one MS1 scan (Orbitrap; 120 K resolution; AGC target 2 × 10 ; max IT 100 ms) followed by data-dependent “Top Speed” (3 s) MS2 scans (collision-induced dissociation; ion trap; NCE 35; AGC 5 × 10³; max IT 100 ms). For multinotch-MS3, the top 10 precursors from each MS2 were fragmented by HCD and analyzed in the Orbitrap (NCE 55; 60 K resolution; AGC 5 × 10 ; max IT 120 ms; 100–500 m/z scan range).

##### Proteomics data analysis

Proteome Discoverer (v2.4; Thermo Fisher) was used for data analysis. MS2 spectra were searched against the UniProt *D. melanogaster* protein database (UP000000803, 2022-11-22) with the following parameters: MS1 and MS2 tolerances of 10 ppm and 0.6 Da, respectively; carbamidomethylation of cysteine (57.02146 Da) and TMT labeling of lysine and peptide N-termini (229.16293 Da) as static modifications; oxidation of methionine (15.9949 Da) and deamidation of asparagine and glutamine (0.98401 Da) as variable modifications. Identified proteins and peptides were filtered at a 1% FDR threshold. Quantitation used high-quality MS3 spectra (average signal-to-noise ratio ≥ 10 and isolation interference < 50%).

### PER-miniTurbo proximity-labeling proteomics analysis

We used a computational pipeline to identify temporally resolved PER-proximal proteins from TMT-multiplexed proximity-labeling proteomics data. The workflow comprises (i) primary mass-spectrometry filtering and isoform collapse, (ii) subcellular-compartment annotation, (iii) differential-abundance significance calling, (iv) a compartment-aware framework that matches each protein to PER-relevant time windows and tests significance at each matched window independently, (v) iterative removal of biologically implausible contaminants, and (vi) functional-category enrichment analysis at both compartment-pooled and per-window resolution.

#### Experimental design and proteomics data input

PER was endogenously tagged with miniTurbo by CRISPR knock-in. Adult fly heads were collected after biotin labeling in one of four 6-h Zeitgeber-time windows (ZT0–6, ZT6–12, ZT12–18, ZT18–24). per-mNeonGreen knock-in flies labeled at ZT0–6 served as a no-ligase control defining the background biotinylation rate. Streptavidin-enriched samples were digested with trypsin, labeled with TMT 10-plex isobaric tags, and analyzed by LC-MS/MS. Protein-level abundances were summarized from peptide spectral matches, yielding 4,654 protein entries, each with TMT-derived log fold-change values and Benjamini–Hochberg-adjusted p-values for every PER-miniTurbo window relative to the per-mNeonGreen control. This table was the input for the pipeline below. The subcellular distribution of PER across the day defines four biologically distinct labeling windows. PER is predominantly nuclear during ZT18–24 (peak concentration after translocation) and ZT0–6 (a declining nuclear pool entering degradation), and predominantly cytoplasmic during ZT6–12 (low cytoplasmic pool) and ZT12–18 (peak cytoplasmic pool, complexed with TIM). These dynamics form the basis of the compartment-aware framework described below.

#### Primary filtering and isoform collapse

The 4,654-row input dataset was filtered using three sequential criteria:

1. Identification confidence: ≥ “High” confidence (Proteome Discoverer FDR-controlled identification at the protein level), rejecting low-confidence single-spectrum hits.
2. Unique peptides: ≥2 unique peptides per protein, excluding single-peptide identifications associated with elevated false-positive rates.
3. Quantification: at least one quantifiable abundance ratio across the four PER-miniTurbo / per-mNeonGreen comparisons.

Redundant accessions for the same gene were then collapsed to a single entry per gene symbol. For genes with multiple accessions, the row with the lowest minimum FDR-adjusted p-value across windows was retained; alternative accessions were logged but excluded from downstream analysis. This yielded a working dataset of 3,255 gene-level proteins.

#### Subcellular compartment annotation

Each protein was assigned a compartment label from a controlled vocabulary: N (nuclear), C (cytoplasmic), N+C (nucleocytoplasmic shuttling), M (membrane or membrane-associated), Mito (mitochondrial), ECM (extracellular matrix), or unannotated. Annotations were assigned in two tiers. First, a regular-expression classifier scanned UniProt and FlyBase description fields for compartment-defining terms, mapping keywords to categories (for example, “ribosomal” → C; “nucleolar”, “histone” and “chromatin” → N), with conflicts (for example, “nuclear envelope” matching both N and M) resolved in favor of the more specific term. Second, for 282 proteins left unassigned, manual curation yielded compartment calls for 230, flagged 52 as contaminants, and retained 7 as unannotated.

#### Significance thresholds

A protein was defined as raw significant at a given window if it satisfied both an enrichment threshold (log fold-change ≥ 0.5, ≈1.4-fold enrichment over the *per*-mNeonGreen control) and a statistical threshold (BH-adjusted p < 0.05). This produces a four-element Boolean vector per protein, independent of compartment annotation. The compartment- and pathway-level enrichments reported in the main text were stable when the enrichment threshold was raised to ≥0.7 and ≥1.0, indicating that the conclusions do not depend on the specific cutoff.

#### Compartment-aware significance assignment

Proximity labeling detects physical closeness, so PER can label a protein only when both occupy the same compartment during a labeling window. We therefore assessed significance only at compartment-matched windows. Nuclear proteins (N) were tested at ZT0–6 and ZT18–24; cytoplasmic and membrane proteins (C, M) at ZT6–12 and ZT12–18; and shuttling (N+C), unannotated, mitochondrial and ECM proteins at all four windows. Within the cleaned dataset of 2,969 proteins, 591 entries were assigned two matched windows and 2,378 were evaluated across all four.

#### Aggregate statistics

The raw-significance criteria (log_2_ FC ≥ 0.5 and FDR-adjusted *p* < 0.05) are evaluated independently at each assigned matched window. A window is designated as “valid significant” (ValidSig) only when it is a matched window and passes both statistical cutoffs; raw significance occurring in non-matched windows is excluded as a likely artifact. This yields a per-window valid-significance containing 0 to *N* True flags (where *N* is the number of matched windows). Complete per-window metrics, including log fold-change, FDR-adjusted *p*-values, raw significance flags, and valid significance flags, are preserved across all four windows for every protein. Proteins may achieve valid significance across zero, one, or multiple matched windows, establishing a multi-timepoint profile for downstream evaluation (e.g., Sirt1 is validly significant only at ZT18–0; CtBP at ZT0–6 and ZT18–0; DnaJ-H across all four windows). Of the 2,969 proteins in the cleaned dataset, 252 were valid significant at one or more matched windows. Of these, 186 (74%) had single-compartment annotations and were tested at two windows (N, 134; C, 33; M, 18), and 66 (26%) were tested at all four windows (N+C, 30; unannotated, 36).

#### Mismatched Set: Proteins Raw-Significant Only at Non-Matched Windows

Proteins that were raw significant at one or more windows but at none of their compartment-matched windows were designated the Mismatched set (n = 30). This set captures compartment misannotations and, potentially, true proximity partners of minority PER sub-pools not reflected in PER’s dominant spatial distribution. Inspection of this set during curation prompted manual reclassification of two proteins to N or N+C and their transfer to the nuclear hit list.

#### Removal of biologically implausible contaminants

After the first significance pass, manual inspection identified four categories of proteins whose enrichment was inconsistent with the cell biology of PER-expressing clock neurons. In each case, high signal in the streptavidin pulldown reflects high abundance in the input rather than proximity to PER.

- Mitochondrial proteins (173 removed). Matrix enzymes, inner-membrane respiratory-chain components, outer-membrane pore proteins and mtDNA-binding factors. PER is not mitochondrial, and this signal class would dominate functional enrichment analysis if retained. This was the largest single removal.
- Yolk and fat-body proteins (52 removed). Yolk proteins (Yp1, Yp2, Yp3) and fat-body metabolic enzymes (Eno, Fum1, fbp, Ahcy, Akr1B, GstO3, Adh, MtnA), which are abundant in adult head extracts owing to fat-body carry-over and are ubiquitous in fly head proteomics datasets.
- Extracellular matrix proteins (41 removed). Collagen subunits, laminins, Pcan, Mp, glutactin and related basement-membrane proteins. Proteins with a single transmembrane domain facing the cytoplasm were retained as M where they had a plausible relationship to clock biology.
- Synaptic proteins (20 removed). Synaptic vesicle proteins (Sap47, nSyb, Nplp2, fax, sif, Esyt2, Nplp1, Syn, Syt1, Syt7, Sytalpha), neurotransmitter-receptor subunits (nAChRalpha4, nAChRalpha6, nAChRbeta1, GluClalpha) and synapse-associated regulators (EndoB, Fas3, Frq1, Sh3beta, mmd)

These removals reduced the working table from 3,255 to 2,969 proteins, of which 252 are valid significant at one or more matched windows. This is the working set used for all downstream analyses.

#### Output lists (Table S1)

- Top_Nuclear (n = 164). Valid significant at ≥1 nuclear-matched window (ZT0–6 or ZT18–24), annotated N or N+C. The primary nuclear proximitome.
- Top_Cytoplasmic (n = 81). Valid significant at ≥1 cytoplasmic-matched window (ZT6– 12 or ZT12–18), annotated C, N+C or M. The primary cytoplasmic proximitome.
- Sig_by_Compartment (n = 252). All valid-significant proteins, listed once each.
- Mismatched (n = 30). Raw significant at one or more windows but at no matched window. Held aside; not used in primary analyses.
- Unknown_Compartment (n = 36). Valid significant at any window but unannotated. Held aside for manual review.
- Hits_ZT0-6 (n = 110), Hits_ZT6-12 (n = 68), Hits_ZT12-18 (n = 65), Hits_ZT18-24 (n = 157). Per-window views of Sig_by_Compartment. A protein may appear in several lists, so these sum to more than 252.

### Clock neuron imaging

Flies were entrained to 12:12 light:dark (LD) cycles for 5–7 days and, where indicated, released into constant darkness (DD) for 6–7 days. Zeitgeber time (ZT) 0 marks the onset of the light phase and ZT12 the onset of the dark phase. Circadian time (CT) refers to times in DD, with CT0 and CT12 corresponding to the times at which the light transitions would have occurred had the LD cycle continued; DD1 and DD2 denote the first and second days in constant darkness.

Flies used for imaging were housed in density-controlled vials (4 females and 4 males) and entrained for 5–7 days. Imaging was performed on 5- to 7-day-old males and females, and no differences in results were observed between sexes. Genotypes and time points are given in the figure legends. Transgenes were expressed in clock neurons using the GAL4/UAS system, with Clk-GAL4 as the pan-clock-neuron driver.

The following combinations were used. PER was imaged using the per-mNeonGreen or per-EGFP knock-in alleles, with Clk-GAL4 > UAS-CD4-tdTomato or UAS-CD8-GFP marking clock-neuron membranes. TIM was imaged using the tim-mNeonGreen knock-in allele. CLK was imaged using the clk-mScarlet-I knock-in allele together with Clk-GAL4 > UAS-CD8-GFP. P-bodies were imaged using the Me31B-Tom2 or Me31B-GFP knock-in alleles, with Clk-GAL4 > UAS-CD8-GFP outlining clock-neuron membranes. For three-color imaging of PER and P-bodies, per-mNeonGreen and Me31B-Tom2 flies were combined with Clk-GAL4 > UAS-BFP-NLS to mark clock-neuron nuclei.

For imaging, 3–4 brains were dissected in chilled Schneider’s Drosophila medium (ThermoFisher Scientific, 21720001) within 5 min under low-light conditions. Punched double-sided tape was used as a spacer on the slide to prevent flattening of the brains. Brains were overlaid with a small volume of ProLong Glass Antifade mounting medium (ThermoFisher Scientific, P36982) and covered with a coverslip. Images were acquired on a Zeiss LSM800 laser-scanning confocal microscope with the Airyscan super-resolution module (125 nm lateral and 350 nm axial resolution), using a 63× Plan-Apochromat oil objective (NA 1.4) and 405, 488, and 561 nm laser lines. Z-stacks (250 nm per slice) or time-lapse series of individual clock neurons were collected and analyzed using Zeiss ZEN software and ImageJ.

### Immunofluorescence staining of brains

Drosophila brains were dissected in chilled Schneider’s Drosophila medium; all brains dissected within a 10-min window were pooled for each experiment. Brains were fixed for 20 min in 4% formaldehyde in PBS at room temperature with gentle rocking, rinsed three times, and permeabilized in PBST (1× PBS, 0.3% Triton X-100) for 1 h at room temperature. Brains were incubated with primary antibody in 5% normal goat serum (NGS) in PBST overnight (at least 16 h) at 4 °C, washed three times for 20 min each in PBST, incubated with secondary antibody in 5% NGS in PBST either overnight at 4 °C or for ∼5 h at room temperature, and washed three times for 20 min each in PBST. Samples were mounted in ProLong Glass Antifade mounting medium (ThermoFisher Scientific, P36982). The following primary antibodies were used: mouse anti-PDF C7 (1:500; Developmental Studies Hybridoma Bank) and mouse anti-V5 (SV5-Pk1, 1:500; Bio-Rad MCA1360). Biotinylated proteins were detected with Alexa Fluor 488-conjugated streptavidin (1:500; ThermoFisher Scientific S11223). Alexa Fluor 488- and Alexa Fluor 555-conjugated secondary antibodies (Molecular Probes) were used at 1:500. Images were acquired on a Zeiss LSM800 laser-scanning confocal microscope with the Airyscan super-resolution detector and a 63× oil-immersion objective and processed using ImageJ.

### HCR-FISH protocol

We adapted a hairpin chain reaction fluorescence in situ hybridization protocol^51^ to detect *per* exonic and *tim* intron 1 transcripts in clock neurons whole-mount brains. Probe sets and amplifier hairpins were designed and synthesized by Molecular Instruments. Flies were dissected in ice-cold PBS, fixed for 20 min in 4% paraformaldehyde at room temperature, and incubated in 70% ethanol overnight at 4 °C. Brains were washed for 5 min at room temperature in wash buffer (2× SSC, 10% formamide, 0.2% Tween-20, 2 mM ribonucleoside vanadyl complex (RVC)), then hybridized overnight at 37 °C in hybridization buffer (2× SSC, 10% dextran sulfate, 10% formamide, 0.2% Tween-20, 2 mM RVC) containing 5 nM probe set. Brains were washed for 1 h in wash buffer and incubated overnight at room temperature in amplification buffer (2× SSC, 10% dextran sulfate, 0.2% Tween-20, 2 mM RVC, 60 nM of each amplifier hairpin). Brains were then washed for 1 h in 2× SSC with 0.2% Tween-20 and mounted in ProLong Glass mountant. For experiments combining HCR-FISH with immunostaining, brains were processed through the full HCR protocol as above, post-fixed in 4% paraformaldehyde for 20 min at room temperature, and then stained with antibodies as described above.

### RNA-seq of FACS-purified clock neurons (*pcm*-RNAi experiments)

*Cell sorting.* Clock neurons were fluorescently labeled using *Clk*-GAL4-driven UAS-GFP-NLS, with DAPI counterstaining to distinguish viable cells. For each replicate, 60–90 brains were dissected and dissociated as previously described^50^. As a dead-cell control, aliquots of dissociated cells were incubated at 60 °C for 5 min to induce DAPI positivity. Live clock neurons were defined by GFP DAPI gating and isolated by fluorescence-activated cell sorting on a Sony MA900. Time from dissection to sorting was approximately 60 min. Approximately 2,000 sorted cells per sample were collected directly into cell lysis buffer (NEBNext Low-Input cDNA Synthesis Module, New England Biolabs) in low-binding 1.5 mL tubes (Eppendorf), giving a final volume of ∼16 µL. Samples were collected at CT0, CT6, CT12 and CT18 from control and *pcm*-RNAi flies, with three biological replicates per condition and time point.

#### Library preparation and sequencing

Following lysis, half of each lysate was processed for cDNA synthesis according to the manufacturer’s protocol. cDNA was assessed by Agilent TapeStation (HSD5000) and Qubit High Sensitivity DNA assay (Fisher Scientific), confirming a mean fragment size of ∼1,500 bp. Libraries were tagmented using the Illumina Nextera XT kit with UDI indexing and reassessed by TapeStation and Qubit, confirming a mean insert size of ∼300 bp. Equimolar pooled libraries, supplemented with 3% PhiX spike-in DNA, were sequenced on an Illumina NextSeq 1000 using 2 × 150 bp paired-end chemistry, targeting 25 M reads per sample.

Adapter sequences were trimmed with TrimGalore (v0.6.7). Reads were aligned to the *Drosophila melanogaster* dm6 genome with transcript annotation from the UCSC Genome Browser using STAR (v2.7.0a) with the parameters --quantMode GeneCounts --outSAMtype BAM SortedByCoordinate. Read coverage was visualized with IGV (v2.16.1).

#### Differential expression and rhythmicity analysis

Gene-level counts from STAR were used for differential expression analysis with DESeq2 (v1.46.0) following standard vignette procedures. Rhythmicity was assessed with JTK_CYCLE across the four time points, searching periods of 20–28 h. Genes were tested and were scored as cycling at BH-adjusted p < 0.05. Of the genes expressed in control samples, 206 met this criterion; the same test was applied to *pcm*-RNAi samples to determine retention or loss of rhythmicity. Because four time points over a single cycle provide limited power for rhythm detection, these calls are conservative and the reported loss of rhythmicity should be interpreted as loss of detectable cycling under this sampling. Oscillation amplitude was calculated as the difference between the maximum and minimum normalized expression values across the four time points. For each gene, peak and trough responses were quantified as the log ratio of *pcm*-RNAi, allowing amplitude changesto be decomposed into peak- and trough-component contributions.

### Clock-neuron-specific ribosome profiling

Ribosome profiling was adapted from a published tissue-specific approach in *Drosophila*^49^. UAS-RpL3-3×FLAG was driven by tim-GAL4 to express FLAG-tagged RpL3 selectively in clock neurons, where it is incorporated into ribosomes. Flies were entrained to 12:12 LD and collected at ZT12. For each replicate, ∼60 brains were dissected in ice-cold PBS containing cycloheximide and flash-frozen; three biological replicates were processed. Brains were homogenized in polysome lysis buffer and lysates cleared by centrifugation, as described previously^49^. An aliquot of each cleared lysate was reserved for total-RNA sequencing as a transcriptional reference, and the remainder was used for ribosome purification. Intact ribosomes were immunoprecipitated with anti-FLAG M2 magnetic beads (Sigma-Aldrich, M8823), and ribosome-protected fragments were generated by RNase I digestion of bead-bound ribosomes. RNA extraction, size selection of ribosome-protected fragments, rRNA depletion and library preparation were performed exactly as described^49^, using the same buffers and commercial kits. Total-RNA libraries were prepared in parallel from the reserved lysate aliquots. Libraries were sequenced on Illumina. Reads were adapter-trimmed with TrimGalore (v0.6.7) and reads mapping to rRNA and tRNA were removed. Remaining reads were aligned to the Drosophila melanogaster dm6 genome (UCSC Genome Browser annotations) using STAR (v2.7.0a). Gene-level counts were used to compute translational efficiency for each gene as the ratio of ribosome-protected fragment reads to total RNA reads^49^.

### rABE Library Preparation and Sequencing

UAS-Me31B-rABE-Flag and UAS-Enc-rABE-Flag flies were crossed to Clk-GAL4 > UAS-GFP-NLS to express the editor fusion proteins together with a nuclear GFP marker in clock neurons. Fusion protein expression was confirmed by anti-FLAG immunostaining. Flies were entrained to 12:12 LD and collected at ZT12. Clock neurons were isolated by fluorescence-activated cell sorting, and libraries were prepared as described in “RNA-seq of FACS-purified clock neurons,” except that sequencing used 2 × 150 bp paired-end chemistry. Three biological replicates of Me31B-rABE, and two Enc-rABE replicates were sequenced.

### rABE bioinformatic processing and quality control

rABE deaminates adenosine to inosine, which is read as guanosine during reverse transcription and therefore appears as an A-to-G mismatch in RNA sequencing, typically at rates of ∼1–10% per site. Editing events were identified following the published analysis pipeline^48^. Reads were trimmed and aligned as described in “RNA-seq bioinformatic processing and data quality control.” Aligned reads were marked for PCR and optical duplicates with Picard MarkDuplicates and split at spliced intron junctions with GATK SplitNCigarReads. Variants were called with bcftools mpileup, generating variant call format files containing variant and reference allele counts at each position. High-confidence editing sites were defined by requiring: ≥20 average reads per site; no detectable A-to-G signal in wild-type controls (allele count ≥2), removing genomic SNPs and endogenous A-to-I edits; significance by Cochran–Mantel–Haenszel test on per-position allele-count contingency tables across replicates, with Benjamini–Hochberg correction (p_adj < 0.05); and recovery in at least two Me31B-rABE replicates. Sites also called in Enc-rABE samples were removed to define ME31B-specific sites.

### Image analysis

A custom ImageJ (Fiji distribution) Python plugin was developed to automate image data analysis. Z-stacks and time-lapse series acquired in Zeiss ZEN software were imported into ImageJ using the Bio-Formats Importer as composite images at lossless 16-bit resolution per channel. Regions of interest (ROIs) were annotated manually, with identical display settings used for all images from the same genotype and neuron type to ensure unbiased comparison.

#### Foci fluorescence intensity

For each cell, a single Z-plane with the largest focus count was selected. Each focus was annotated with a polygon ROI, and mean pixel brightness (arbitrary units, a.u.) and area (µm²) were obtained using built-in ImageJ functions. Integrated intensity was calculated as the product of mean brightness and area (a.u. × µm²). Background fluorescence, estimated as the mean intensity of a background selection in the same plane, was subtracted from the integrated intensity of each ROI.

#### Foci counts

Foci were annotated manually where clearly distinguishable from background and counted across all Z-planes for each cell. This procedure was used to quantify PER nuclear foci, CLK nuclear foci and ME31B cytoplasmic puncta.

#### Three-dimensional segmentation of P-bodies

ME31B puncta were segmented in three dimensions from Z-stacks using Imaris 11.0.1 software. From the maximum Z projection, the lLNVs were demarcated using the ‘Surface creation’ feature with an assigned diameter of 6 uM. Me31B protein puncta and *per* exons were demarcated using the ‘Spots creation’ feature with an assigned diameter of 0.5 µm and a ‘distance from surface’ value between -20 to -0.5 uM. *pe*r exons were divided into two categories: those within 0.4 µm of the nearest Me31B puncta, and those within 0.4 µm to 1 µm of the nearest Me31B puncta. Videos and snapshots of the 3D projections were generated in Imaris 11.0.1.

#### Association of mRNA with P-bodies

*per* and *tim* mRNA foci (HCR-FISH), PER-mNG foci and ME31B puncta were segmented independently in three dimensions as above. An mRNA or PER focus was scored as P-body-associated when its centroid lay within 0.4 µm of the nearest ME31B object centroid. The associated fraction was calculated per cell as the number of associated foci divided by the total number of foci of that species in the cell. For the distance analysis shown in Figure 4E, the centroid-to-centroid distance from each *per* mRNA focus to the nearest ME31B object was computed.

#### Adjacency of PER foci to P-bodies

PER-mNG foci and Me31B-Tom2 puncta were segmented independently in three dimensions. A PER focus was scored as adjacent to a P-body when its centroid lay within 0.4 µm of the nearest ME31B object centroid, the same criterion used for mRNA association. The adjacent fraction was calculated per cell as the number of adjacent PER foci divided by the total number of PER foci in that cell.

### HCR-FISH data analysis

We followed a previously established protocol for analysis of FISH data, involving segmentation of clock neurons and detection of diffraction-limited spots^61^. Clock neurons were delineated by the Clk>CD8-GFP membrane marker, and spot counts per neuron were obtained from the segmented volumes. To assess association between per or tim mRNA and P-bodies, we computed all pairwise distances between spots detected in the two channels within the same neuron. For each mRNA spot, the nearest-neighbor distance (NND) was defined as the smallest three-dimensional distance to any ME31B spot in that neuron. Spots with an NND below 400 nm were classified as P-body-associated, allowing the associated fraction to be calculated per cell as the number of associated spots divided by the total number of mRNA spots. The same criterion was applied to quantify the association of PER-mNG foci with ME31B puncta. The custom ImageJ plugin for ROI-based spot classification is available on GitHub (https://github.com/yeyuan98/punctaTracker) and via an ImageJ update site (https://sites.imagej.net/Yuanye1998/). The workflow for downstream processing of classified spots and nearest-neighbor distance computation is documented in the R package ijAnalysis (https://yeyuan98.r-universe.dev/articles/ijAnalysis/spotInRoi.html).

### Locomotor activity and rhythmicity analysis

Individual adult male flies (5-7 days old) were placed in glass capillary tubes (∼4 mm inner diameter, 5 cm in length) containing 2% agar and 4% sucrose food, which were then loaded into TriKinetics DAM2 Drosophila Activity Monitors (Waltham, MA, USA) for locomotor activity recordings. Flies were entrained to Light-Dark (LD) cycles with lights on for 12 hours and off for 12 hours for ∼5 days, followed by complete darkness (DD) for ∼7 days. Beam crossing counts were placed into 30-minute bins for time-series analysis of locomotor activity. Averaged population activity profiles under Light-Dark cycles and constant conditions were generated using a commercially available software ClockLab (Actimetrics) and public domain R Rethomics software package. Activity of individual flies under complete darkness conditions following LD cycles was used to analyze rhythmicity and determine the free-running period of the circadian clock. Rhythmicity and free-running period of individual flies were determined by a chi-square periodogram analysis with a confidence level of 0.001 using the ClockLab software. The “Power” and “Significance” values generated from the chi-square analysis were used to calculate “Rhythmic Power” as a measure of the strength of each rhythm.

### Statistical analysis

For protein fluorescence and RNA spot-count measurements, data were pooled from hemi-brain images collected across at least three independent experiments. Measurements were taken from distinct neurons within a brain, and no neuron was measured more than once. Sequencing experiments used two to three biological replicates with ∼60 flies per replicate, and behavioral experiments used ∼30 flies per genotype. Live imaging and biochemical experiments used ∼7-day-old males and females; behavioral experiments used males only, except for the *per ¹* rescue, in which females were used. All statistical analyses were performed in GraphPad Prism 11. Comparisons between two groups used Student’s *t*-test with Welch’s correction; multiple comparisons used ANOVA with Brown–Forsythe and Welch’s corrections. Significance is indicated as ns (not significant), * p < 0.05, ** p < 0.01, *** p < 0.001, **** p < 0.0001. Error bars represent mean ± SEM. Sample sizes, statistical tests and exact p-values for all experiments are given in Table S3.

## DATA REPOSITORY

Sequencing data from all the experiments are deposited in GSE301091 and GSE300438 repositories.

## Supplementary tables

**Table S1: TMT mass spectrometry analysis.** List of all PER-proximal proteins, significant PER-proximal proteins at ZT0-6, ZT6-12, ZT12-18, ZT18-0, compartment-specific significant PER-proximal proteins from *per*-V5-miniturbo *vs. per*-mNG TMT mass spectrometry analysis.

**Table S2: Locomotor behavior data.** Genotypes, no. of flies, percent rhythmicity, power, and period values of all locomotor behavior experiments.

**Table S3: Statistical significance.** Adjusted P values between time points for all experiments. ns: non-significant, * <0.05, ** <0.01, *** <0.001, **** <0.0001.

## ACKNOWLEDGEMENTS

We thank Venkatesh Basrur for help with the design of proximity labeling and mass-spec experiments. We thank Elizabeth Gavis and Florence Besse for providing the Me31B-GFP and Me31-tdTom flylines. We thank Dunham Clark and Brelian Khatib-Shahidi for help with initial set of experiments. We thank the Bloomington Drosophila Stock Center for providing fly strains. This work was supported by National Institutes of Health NIGMS R35GM133737 grant to S.Y., the Alfred P. Sloan fellowship, the McKnight scholar grant, and the Chan Zuckerberg collaborative pairs grant to S.Y., and the Rackham Predoctoral Fellowship to Y.Y..

## AUTHOR CONTRIBUTIONS

Y.X. performed the miniTurbo experiments. Y.X., S.C., and D.B. performed all the imaging experiments. Y.X. and Y.Y. performed the sequencing experiments. S.Y. designed the project and experimental plan. Y.X., S.C., and S.Y. wrote the manuscript with input from all authors.

## CONFLICTS OF INTEREST

The authors declare no financial conflict of interest.

## Extended Data Figures

**Extended Data Figure 1.**
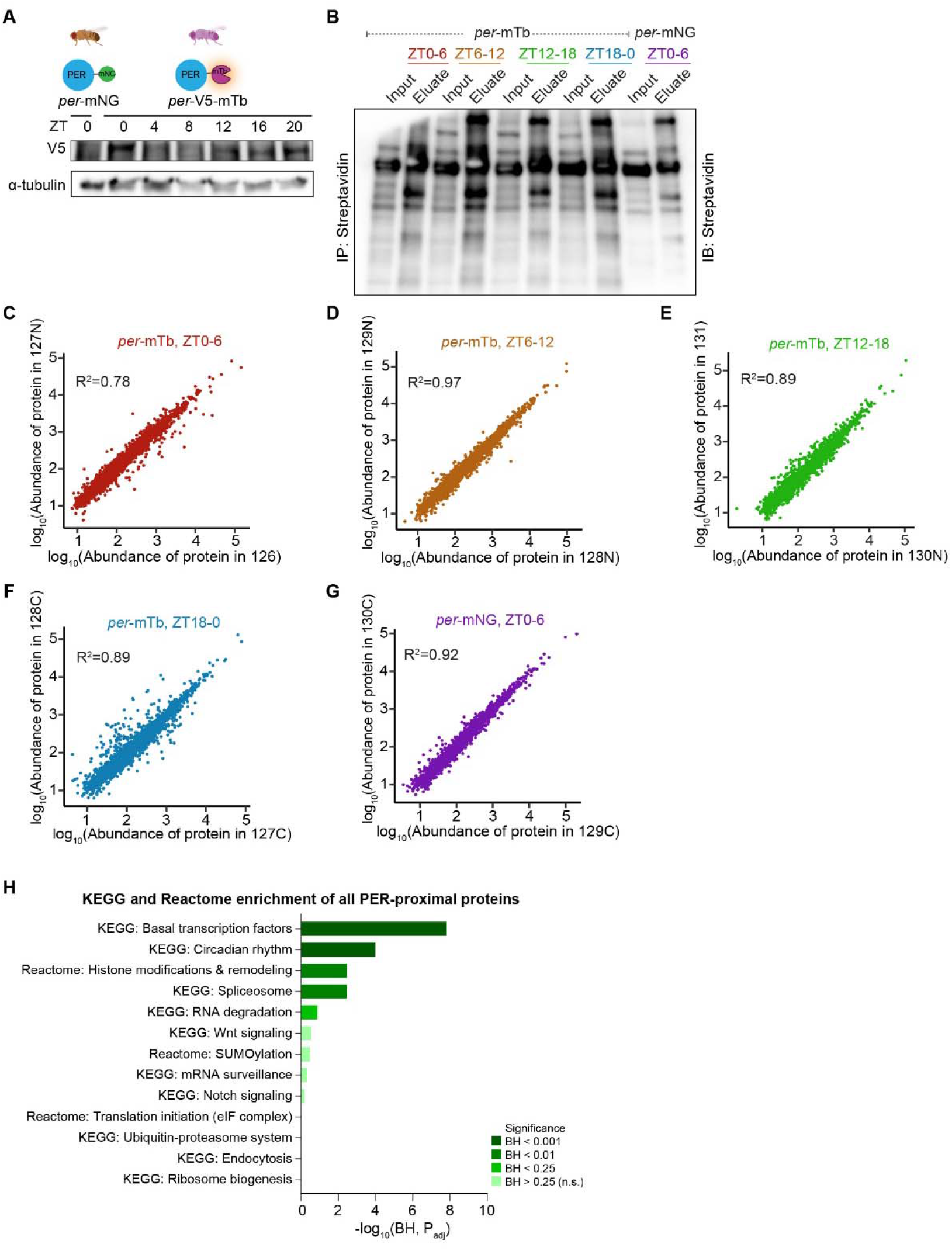
Validation of the PER-V5-miniTurbo knock-in and quality control of the proximity-labeling proteomics. **(A)** Anti-V5 immunoblot of head lysates from *per*-V5-miniTurbo flies collected every 4 h (ZT0–ZT20) and from *per*-mNeonGreen flies at ZT0. α-Tubulin, loading control. **(B)** Streptavidin blot of input and eluate fractions from *per*-V5-miniTurbo (ZT0–6, ZT6–12, ZT12–18, ZT18–24) and *per*-mNeonGreen (ZT0–6) head lysates following streptavidin pull-down. **(C–G)** Correlation between biological replicates for *per*-V5-miniTurbo at ZT0–6 (C), ZT6–12 (D), ZT12–18 (E) and ZT18–24 (F), and for *per*-mNeonGreen at ZT0–6 (G). Axes show log protein abundance in the two TMT channels assigned to each replicate pair; R² is given in each panel. **(H)** KEGG and Reactome enrichment of all PER-proximal proteins (n = 252). Bars are shaded by Benjamini–Hochberg-adjusted significance. mTb, miniTurbo; mNG, mNeonGreen; IP, immunoprecipitation; IB, immunoblot; ZT, Zeitgeber time; BH, Benjamini–Hochberg.

**Extended Data Figure 2.**
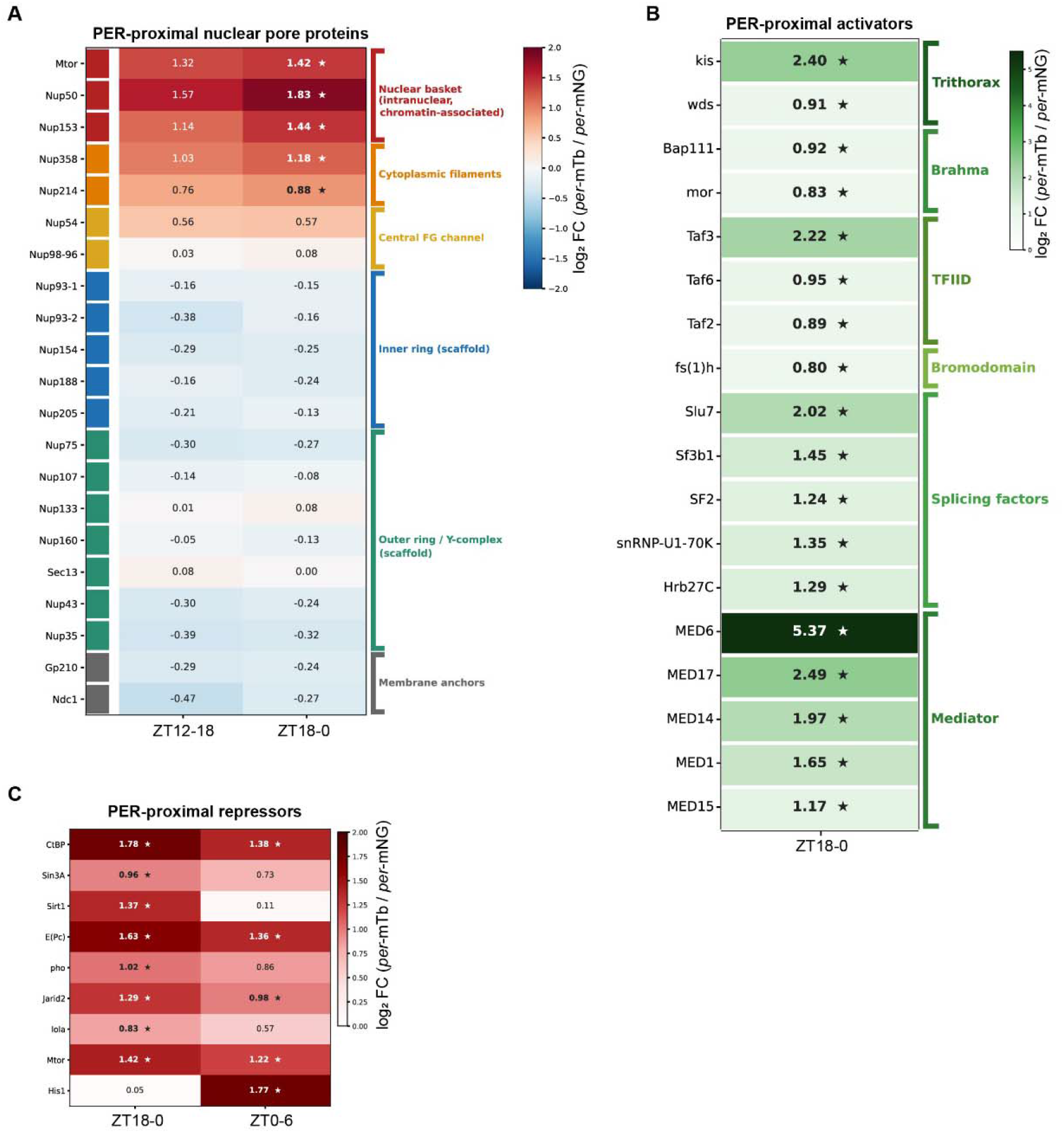
PER’s nuclear neighborhood distinguishes phase-restricted association with transcriptional machinery from sustained association with chromatin silencers. **(A)** Heat map of log fold change (*per*-V5-mTb / *per*-mNeonGreen) for nuclear pore complex components at ZT12–18 and ZT18–24. Nucleoporins are grouped by their position within the pore (nuclear basket, cytoplasmic filaments, central FG channel, inner and outer scaffold rings, membrane anchors). Enrichment is restricted to basket and cytoplasmic-filament nucleoporins; scaffold and membrane-anchor components are not enriched. **(B)** Log fold change for PER-proximal transcriptional activators and coactivators at ZT18–24, grouped by complex (Trithorax, Brahma, TFIID, bromodomain, splicing factors, Mediator). **(C)** Log fold change for PER-proximal transcriptional repressors at ZT18–24 and ZT0–6. Asterisks indicate proteins meeting the proximity criteria (log FC ≥ 0.5, FDR-adjusted p < 0.05) at that window. FC, fold change; mTb, miniTurbo; mNG, mNeonGreen; ZT, Zeitgeber time.

**Extended Data Figure 3.**
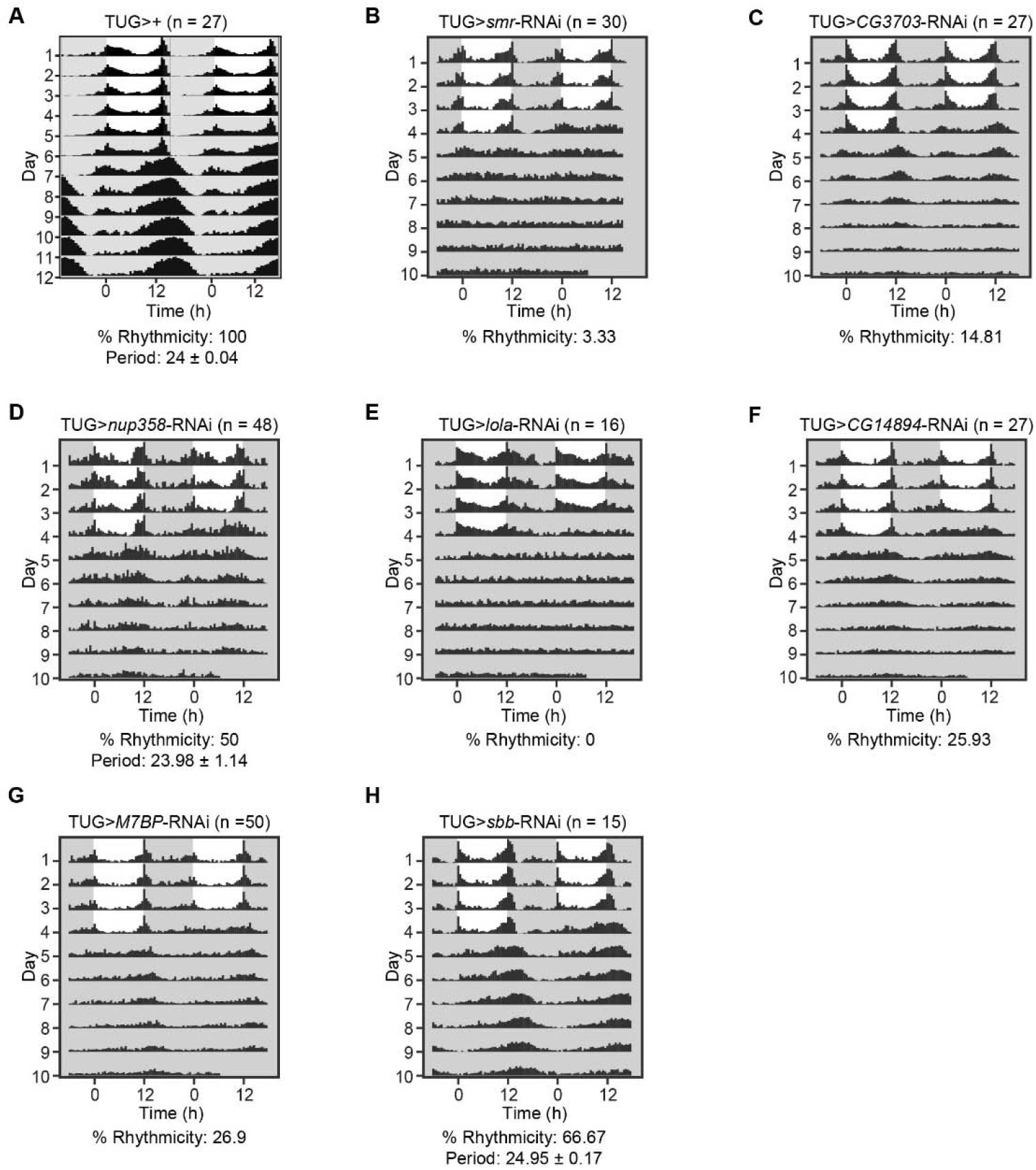
Locomotor rhythms of additional RNAi lines from the PER-proximal screen. (A–H) Flies were entrained to 12:12 LD (ZT0, lights on; ZT12, lights off) for 4 days and released into constant darkness for 6 days. Double-plotted population actograms are shown for *tim*-UAS-GAL4 (TUG) > + (n = 27) (A), TUG > *smr*-RNAi (n = 30) (B), TUG > *CG3703*-RNAi (n = 27) (C), TUG > *nup358*-RNAi (n = 48) (D), TUG > *lola*-RNAi (n = 16) (E), TUG > *CG14894*-RNAi (n = 27) (F), TUG > *M7BP*-RNAi (n = 50) (G) and TUG > *sbb*-RNAi (n = 15) (H). Percent rhythmicity is shown below each panel, with free-running period (mean ± SEM) given where enough flies were rhythmic to determine it. TUG, *tim*-UAS-GAL4; ZT, Zeitgeber time; LD, light:dark; DD, constant darkness.

**Extended Data Figure 4.**
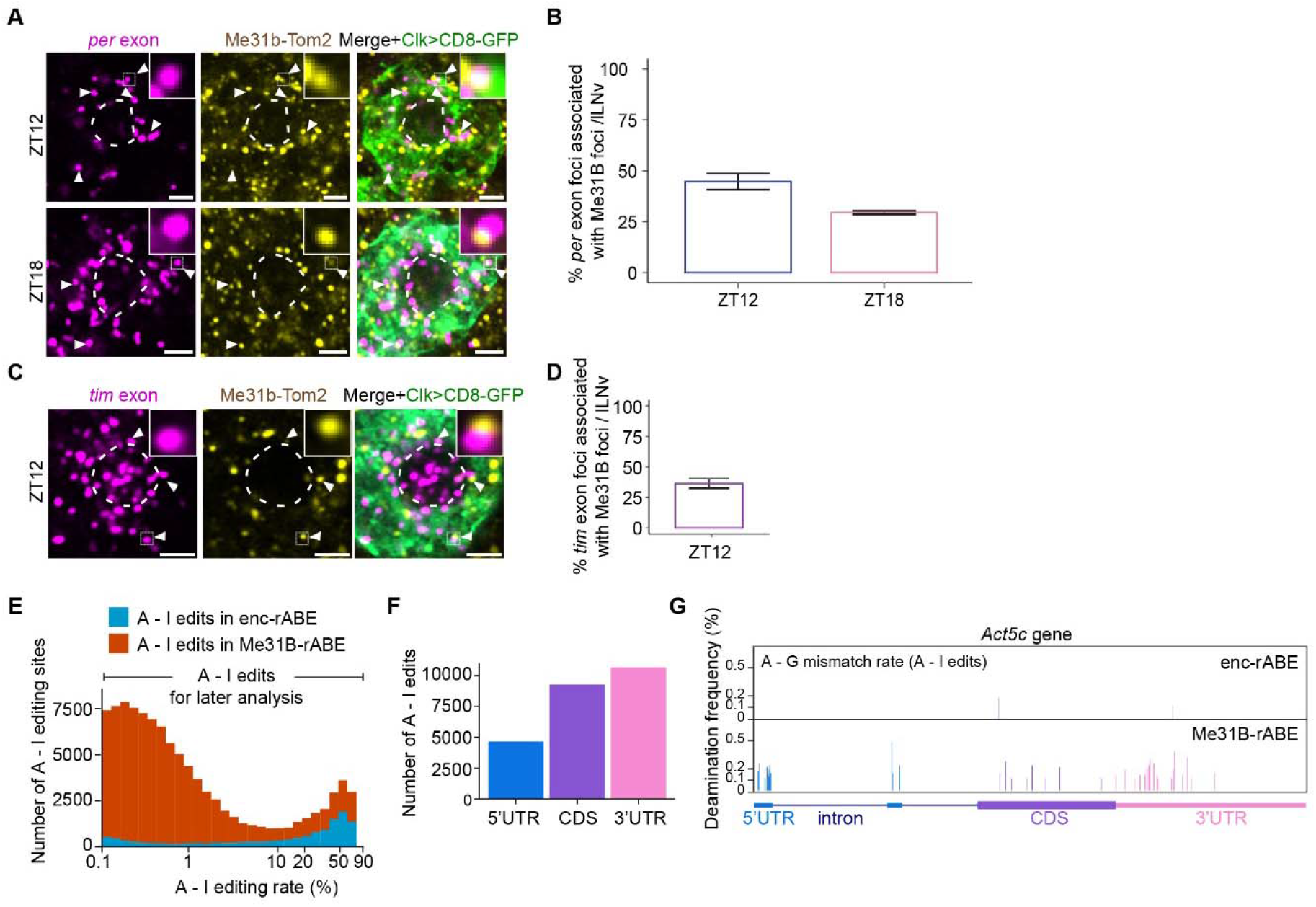
Association of *per* and *tim* mRNAs with P-bodies in an independent Me31B knock-in line, and characterization of Me31B-rABE editing. **(A)** Representative images of *per* exonic HCR-FISH foci (magenta) and Me31B-Tom2 (yellow) in large ventral lateral neurons (lLNv) of *Clk*>CD8-GFP; Me31B-Tom2 flies at ZT12 and ZT18. Membranes are marked by CD8-GFP (green); arrowheads mark *per* foci associated with Me31B foci, and insets show magnified examples. Dashed lines demarcate nuclei. Scale bars, 5 µm. **(B)** Percentage of *per* foci associated with Me31B foci per lLNv at ZT12 and ZT18, quantified from (A). n = 24 neurons per time point. **(C)** Representative images of *tim* exonic HCR-FISH foci (magenta) and Me31B-Tom2 (yellow) in lLNv of the same genotype at ZT12, labeled as in (A). Scale bars, 5 µm. **(D)** Percentage of *tim* foci associated with Me31B foci per lLNv at ZT12, quantified from (C). n = 17 neurons per time point. **(E)** Distribution of A-to-I editing rates across all called sites in Me31B-rABE (orange) and Enc-rABE (blue) samples. Sites in the low-rate range were retained for downstream analysis; the high-rate tail is dominated by sites shared with controls, consistent with endogenous ADAR activity. **(F)** Distribution of Me31B-rABE-specific editing sites across 5′UTR, CDS and 3′UTR regions. **(G)** Gene-track view of A-to-I editing along *Act5C* in Enc-rABE and Me31B-rABE samples. ZT, Zeitgeber time; lLNv, large ventral lateral neuron.

**Extended Data Figure 5.**
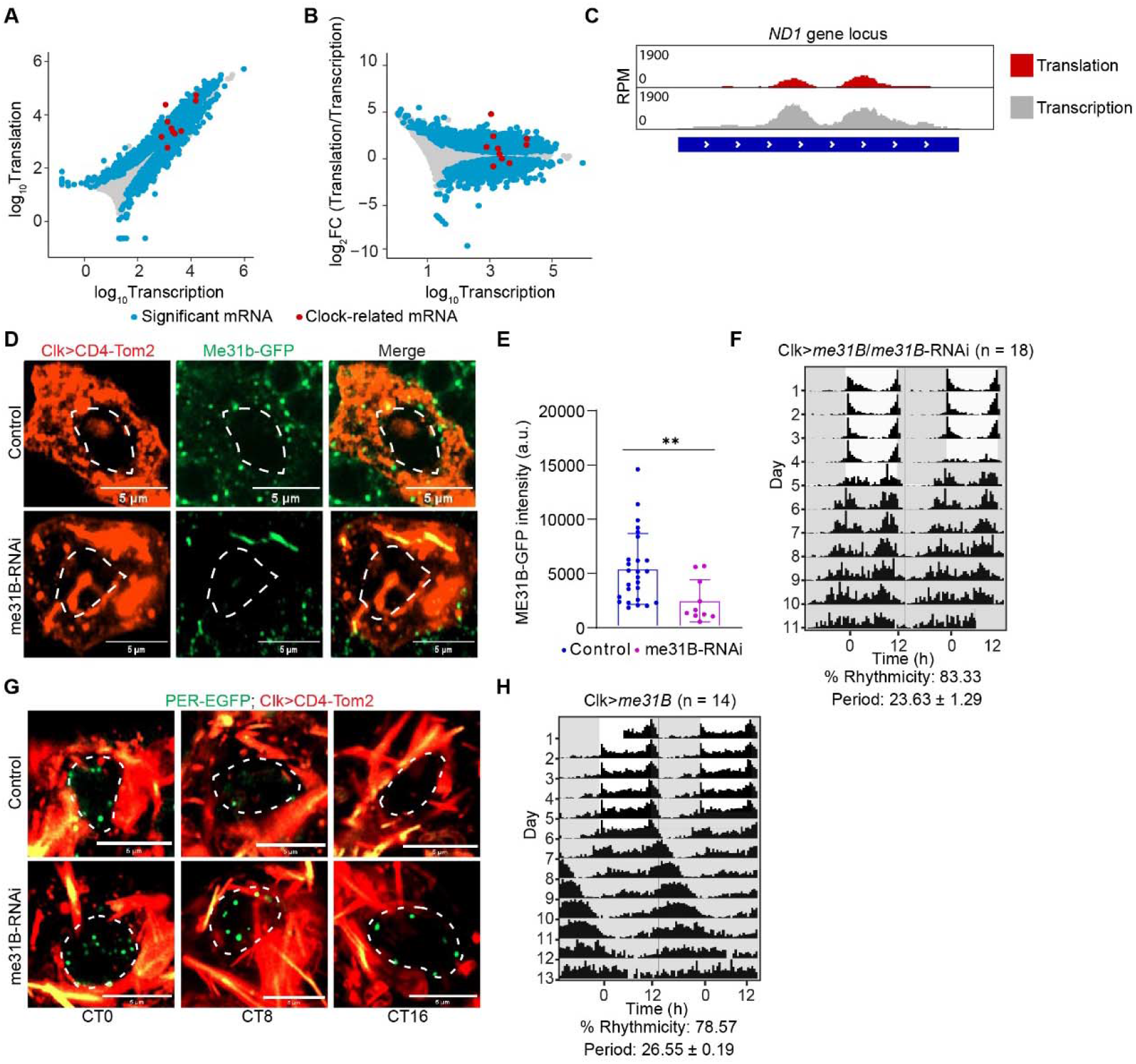
Clock-neuron translatome controls and validation of Me31B knockdown and overexpression. **(A)** Translation versus transcription for the clock-neuron translatome at ZT12 (log reads). Blue, significantly changed transcripts; red, clock-related transcripts. **(B)** Translational efficiency, plotted as log (translation/transcription), against transcription at ZT12. Colors as in (A). **(C)** Gene-track view of ribosome-protected fragment reads (translation, red) and total RNA reads (transcription, grey) at the *ND1* locus, shown as an abundant transcript with low translational enrichment. RPM, reads per million mapped reads. **(D)** Representative images of Me31B-GFP (green) in large ventral lateral neurons (lLNv) of *Clk*>CD4-tdTomato (control, top) and *Clk*>CD4-tdTomato; *me31B*-RNAi (bottom) flies. Membranes are marked by CD4-tdTomato (red); dashed lines demarcate nuclei. Note the loss of discrete Me31B puncta in the knockdown. Scale bars, 5 µm. **(E)** Me31B-GFP intensity per lLNv in control and *me31B*-RNAi flies, quantified from (D). **(F)** Double-plotted population actogram of *Clk*>UAS-Me31B/*me31B*-RNAi rescue flies (n = 18) entrained to 12:12 LD and released into constant darkness. Percent rhythmicity and free-running period (mean ± SEM) are shown below. **(G)** Representative images of PER-EGFP foci (green) in lLNv of *per*-EGFP; *Clk*>CD4-tdTomato (control) and *per*-EGFP; *Clk*>CD4-tdTomato; *me31B*-RNAi flies at CT0, CT8 and CT16 in constant darkness; quantified in Figure 5F. Scale bars, 5 µm. **(H)** Double-plotted population actogram of *Clk*>UAS-Me31B overexpression flies (n = 14), plotted as in (F). lLNv, large ventral lateral neuron; FC, fold change; RPM, reads per million mapped reads; CT, circadian time; ZT, Zeitgeber time.

## Notes

### Competing Interest Statement

The authors have declared no competing interest.

